# Female-specific upregulation of insulin pathway activity mediates the sex difference in *Drosophila* body size plasticity

**DOI:** 10.1101/2020.04.22.054239

**Authors:** Jason W. Millington, Chien Chao, Ziwei Sun, Paige J. Basner-Collins, George P. Brownrigg, Lianna W. Wat, Bruno Hudry, Irene Miguel-Aliaga, Elizabeth J. Rideout

**Author notes:** Corresponding author: Department of Cellular and Physiological Sciences, Life Sciences Institute, The University of British Columbia, 2350 Health Sciences Mall, Vancouver, BC, V6T 1Z3, Canada.

## Abstract

Nutrient-dependent body size plasticity differs between the sexes in most species, including mammals. Previous work in *Drosophila* showed that body size plasticity was higher in females, yet the mechanisms underlying the sex difference in body size plasticity remain unclear. Here, we discover that a protein-rich diet augments body size in females and not males because of a female-specific increase in activity of the conserved insulin/insulin-like growth factor signaling pathway (IIS). This increased IIS activity was triggered by a diet-induced increase in *stunted*, and required *Drosophila insulin-like peptide 2*, illuminating new sex-specific roles for these genes. Importantly, we show that sex determination gene *transformer* regulates the diet-induced increase in *stunted* and IIS activity, and mediates the sex difference in body size plasticity. This identifies one sex-specific mechanism underlying the nutrient-dependent regulation of IIS activity and body size plasticity, providing vital insight into conserved mechanisms that mediate sex differences in phenotypic plasticity.

## INTRODUCTION

In insects, as in many animals, the rate of growth during development is influenced by environmental factors such as nutrient availability (Boulan et al., 2015; Edgar, 2006; Hietakangas & Cohen, 2009; Nijhout, 2003; Nijhout et al., 2014). When nutrients are abundant, the rate of growth is high and body size is large (Beadle et al., 1938; Edgar, 2006; Mirth & Shingleton, 2012; Nijhout, 2003; Robertson, 1963). When nutrients are scarce, the rate of growth is lower and body size is smaller (Beadle et al., 1938; Edgar, 2006; Mirth & Riddiford, 2007; Mirth & Shingleton, 2012; Nijhout, 2003; Robertson, 1963). This ability of an organism or genotype to adjust its body size in line with nutrient availability is a form of phenotypic plasticity (Agrawal, 2001; Garland & Kelly, 2006). While the capacity of individuals to display nutrient-dependent changes to body size depends on many factors, one important factor that affects phenotypic plasticity is whether an animal is male or female (Stillwell et al., 2010; Teder & Tammaru, 2005). For example, studies in *Drosophila* found that the magnitude of changes to wing cell size and cell number in a nutrient-poor diet were larger in females compared with males (Alpatov, 1930). Additionally, recent studies that systematically manipulated dietary carbohydrates and protein confirmed that the magnitude of diet-induced changes to some morphological traits was larger in female flies (Shingleton et al., 2017). While these studies clearly establish a sex difference in nutrient-dependent phenotypic plasticity, the genetic and molecular mechanisms underlying this increased trait size plasticity in females remain unclear.

Clues into the genes and pathways that may underlie the increased nutrient-dependent phenotypic plasticity in female flies have emerged from over 20 years of studies on nutrient-dependent growth in *Drosophila* (Andersen et al., 2013; Boulan et al., 2015; Edgar, 2006; Koyama & Mirth, 2018; Mirth & Piper, 2017). In particular, these studies have identified the conserved insulin/insulin-like growth factor signaling pathway (IIS) as a key regulator of nutrient-dependent growth in *Drosophila* (Böhni et al., 1999; Britton et al., 2002; Chen et al., 1996; Fernandez et al., 1995; Grewal, 2009; Teleman, 2009). In nutrient-rich conditions, insulin-producing cells (IPCs) in the larval brain release *Drosophila* insulin-like peptides (dILPs) into the circulation (Brogiolo et al., 2001; Géminard et al., 2009; Ikeya et al., 2002; Rulifson et al., 2002). These dILPs bind the Insulin-like Receptor (InR; FBgn0283499) on target cells to induce receptor autophosphorylation and recruitment of adapter proteins such as InR substrate Chico (*chico*; FBgn0024248) and Lnk (*Lnk*; FBgn0028717) (Almudi et al., 2013; Böhni et al., 1999; Chen et al., 1996; Poltilove et al., 2000; Werz et al., 2009). These adapter proteins, when phosphorylated, enable the recruitment of a regulatory subunit of the *Drosophila* homolog of phosphatidylinositol 3-kinase (*Pi3K21B*; FBgn0020622) that recruits and activates the catalytic subunit of Pi3K (*Pi3K92E*; FBgn0015279). This activated Pi3K complex catalyzes the production of phosphatidylinositol (3,4,5)- trisphosphate (PIP_3_) from phosphatidylinositol (4,5)-bisphosphate (PIP_2_) (Leevers et al., 1996). The increased abundance of PIP_3_ in the plasma membrane recruits and activates signaling proteins such as phosphoinositide-dependent kinase 1 (Pdk1; FBgn0020386) and Akt (Akt; FBgn0010379), which influence diverse cellular processes to enhance cell, tissue, and organismal growth (Cho et al., 2001; Grewal, 2009; Rintelen et al., 2001; Verdu et al., 1999). On the other hand, in nutrient-restricted conditions, dILP release from the IPCs is reduced (Géminard et al., 2009), and plasma membrane Pi3K recruitment, PIP_3_ levels, and Pdk1- and Akt-dependent signaling are all reduced (Britton et al., 2002; Nowak et al., 2013). Together, these changes diminish cell, tissue, and organismal growth (Arquier et al., 2008; Britton et al., 2002; Géminard et al., 2009; Honegger et al., 2008; Okamoto et al., 2013; Rulifson et al., 2002; Zhang et al., 2009). Indeed, the potent growth-promoting ability of IIS activation is demonstrated by studies in *Drosophila* showing that genetic manipulations that increase IIS activity augment growth during development (Arquier et al., 2008; Goberdhan et al., 1999; Honegger et al., 2008; Ikeya et al., 2002; Nowak et al., 2013; Okamoto et al., 2013; Oldham et al., 2002), whereas genetic mutations that lower IIS activity strongly reduce cell, organ, and body size (Böhni et al., 1999; Brogiolo et al., 2001; Chen et al., 1996; Colombani et al., 2003; Gao et al., 2000; Grönke et al., 2010; Leevers et al., 1996; Murillo-Maldonado et al., 2011; Rulifson et al., 2002; Weinkove et al., 1999; Zhang et al., 2009). Because increased IIS activity is sufficient to bypass the reduced cell growth normally observed upon nutrient restriction (Britton et al., 2002; Géminard et al., 2009; Nowak et al., 2013), and that mutations blunting IIS pathway activity decrease growth even in nutrient-rich conditions (Böhni et al., 1999; Brogiolo et al., 2001; Chen et al., 1996; Leevers et al., 1996), studies in *Drosophila* have established a role for IIS in promoting organismal growth downstream of nutrient input. While this highlights the vital role that *Drosophila* studies have played in elucidating the mechanisms by which IIS couples nutrient input with cell, tissue, and organismal growth, it is important to note that most studies in this area used a mixed-sex population of larvae. Given that cell and body size are significantly different between male and female flies (Alpatov, 1930; Brown & King, 1961; Okamoto et al., 2013; Partridge et al., 1994; Rideout et al., 2015; Sawala & Gould, 2017; Testa et al., 2013), more knowledge is needed of nutrient-dependent changes to body size and IIS activity in each sex.

Recent studies have begun to make progress in this area by studying IIS regulation and function in both sexes in a single dietary context (reviewed in Millington & Rideout, 2018). One study on late third instar larvae reported sex differences in *dilp* mRNA levels, in IIS activity, and in the release of dILP2, an important growth-promoting dILP, from the IPCs (Rideout et al., 2015). Similarly, transcriptomic studies have detected male-female differences in mRNA levels of genes associated with IIS function (Mathews et al., 2017; Rideout et al., 2015), and revealed links between IIS and the sex determination hierarchy gene regulatory network (Castellanos et al., 2013; Chang et al., 2011; Clough et al., 2014; Fear et al., 2015; Garner et al., 2018; Goldman & Arbeitman, 2007). As evidence of sex-specific IIS regulation continues to accumulate, several reports have also identified sex-limited and sex-biased phenotypic effects caused by changes to IIS function. For example, changes to IIS activity show sex-biased effects on larval growth and final body size (Grönke et al., 2010; Rideout et al., 2015; Shingleton et al., 2005; Testa et al., 2013). In adults, widespread sex-specific and sex-biased changes to gene expression were observed in flies with altered diet and IIS activity (Camus et al., 2019; Graze et al., 2018). Further, sex differences exist in how changes to diet and IIS activity affect life span (Bjedov et al., 2010; Clancy et al., 2001; Giannakou et al., 2004; Grönke et al., 2010; Regan et al., 2016; Tatar et al., 2001; Woodling et al., 2020; Wu et al., 2020). Together, these studies illuminate the utility of *Drosophila* in revealing sex-specific IIS regulation and describing the physiological impact of this regulation. Yet, more studies are needed to discover the molecular mechanisms underlying sex-specific IIS regulation, and to extend these studies beyond a single nutritional context.

Additional insights into potential mechanisms underlying the sex difference in nutrient-dependent trait size plasticity come from studies on the regulation of cell, tissue, and body growth by sex determination genes. In flies, sex is determined by the number of X chromosomes. In XX females, a functional splicing factor called Sex-lethal (Sxl; FBgn0264270) is produced (Bell et al., 1988; Bridges, 1921; Cline, 1978; Salz & Erickson, 2010). Sxl protein binds to the pre-mRNA of *transformer* (*tra*, FBgn0003741), Sxl’s most well-known target gene, where the Sxl-dependent splicing of *tra* pre-mRNA allows a functional Tra protein to be produced in females (Belote et al., 1989; Boggs et al., 1987; Inoue et al., 1990; Sosnowski et al., 1989). In XY males, no functional Sxl protein is produced (Cline & Meyer, 1996; Salz & Erickson, 2010). As a result, *tra* pre-mRNA undergoes default splicing, and no functional Tra protein is produced in males (Belote et al., 1989; Boggs et al., 1987; Inoue et al., 1990; Sosnowski et al., 1989). An extensive literature shows that the presence of functional Sxl and Tra proteins in females accounts for most aspects of female sexual development, behavior, and physiology (Anand et al., 2001; Billeter et al., 2006; Brown & King, 1961; Camara et al., 2008; Christiansen et al., 2002; Clough et al., 2014; Dauwalder, 2011; Demir & Dickson, 2005; Goodwin et al., 2000; Hoshijima et al., 1991; Hudry et al., 2016, 2019; Ito et al., 1996; Millington & Rideout, 2018; Neville et al., 2014; Nojima et al., 2014; Pavlou et al., 2016; Pomatto et al., 2017; Regan et al., 2016; Rezával et al., 2014, 2016; Rideout et al., 2010; Ryner et al., 1996; Sturtevant, 1945; von Philipsborn et al., 2014). Recently, roles for Sxl and Tra in regulating the sex difference in body size were also described. While *Drosophila* females are normally significantly and visibly larger than male flies, females lacking neuronal *Sxl* are significantly smaller than control females, and no longer different in size from males (Sawala & Gould, 2017). Interestingly, *Sxl* function in specific neurons, the IPCs and GAD1-GAL4-positive neurons, mediate its effects on female growth during development (Sawala & Gould, 2017). Similarly, females lacking a functional Tra protein were significantly smaller than control females; however, these *tra* mutant females were still larger than males (Brown & King, 1961; Mathews et al., 2017; Rideout et al., 2015). Together, these studies indicate a requirement for both Tra and Sxl in promoting a larger body size in females, providing vital insight into the intersection between the sex determination pathway and the regulation of body size. However, much remains to be discovered about the mechanisms by which Sxl and Tra impact body size. Moreover, it remains unclear whether sex determination genes contribute to the male-female difference in diet-induced trait size plasticity, as previous studies on sex determination genes used a single nutritional context.

In the present study, we aimed to improve knowledge of the genetic and molecular mechanisms that contribute to male-female differences in nutrient-dependent plasticity in *Drosophila.* Our detailed examination of nutrient-dependent body size plasticity revealed increased phenotypic plasticity in females in response to a protein-rich diet, in line with prior studies on trait size plasticity (Shingleton et al., 2017). Importantly, we show that a nutrient-dependent upregulation of IIS activity in females and not in males in a protein-rich context is responsible for the increased body size plasticity in females. Mechanistically, we show that a nutrient-dependent upregulation of *stunted* (*sun*; FBgn0014391) mRNA levels in females triggers the diet-induced increase in IIS activity, as females lacking *sun* do not augment IIS activity or body size in a protein-rich diet. Importantly, we show that sex determination gene *tra* is required for the nutrient-dependent increase in *sun* mRNA, IIS activity, and phenotypic plasticity in females, and that ectopic *tra* expression in males enhances nutrient-dependent body size plasticity via *sun*-mediated regulation of IIS activity. Together, these results provide new insight into the molecular mechanisms that govern male-female differences in body size plasticity, and identify a previously unrecognized role for sex determination gene *tra* in regulating nutrient-dependent phenotypic plasticity.

## RESULTS

### High levels of dietary protein are required for increased nutrient-dependent body size plasticity in females

Previous studies identified a sex difference in nutrient-dependent plasticity in several morphological traits (Shingleton et al., 2017; Stillwell et al., 2010; Teder & Tammaru, 2005). To determine whether sex differences in nutrient-dependent body size plasticity exist in *Drosophila*, we measured pupal volume, an established readout for *Drosophila* body size (Delanoue et al., 2010), in *white^1118^* (*w*; FBgn0003996) males and females reared on diets of varying nutrient quantity. We found that pupal volume in *w^1118^* female larvae raised on the 2-acid diet (1×) (Lewis, 1960) was significantly larger than genotype-matched females raised on a diet with half the nutrient quantity (0.5×) (Fig. 1A). In *w^1118^* males, pupal volume was also significantly larger in larvae raised on the 1×diet compared with the 0.5× diet (Fig. 1A). No significant sex-by-diet interaction was detected using a two-way analysis of variance (ANOVA) (sex:diet interaction *p* = 0.7048; S1 Table), suggesting that nutrient-dependent body size plasticity was not different between the sexes in this context. We next compared pupal volume in *w^1118^* males and females raised on the 1× diet with larvae cultured on a diet with twice the nutrient content (2×). Pupal volume in *w^1118^* females was significantly larger in larvae raised on the 2× diet compared with larvae cultured on the 1× diet (Fig. 1A). In *w^1118^* males, the magnitude of the nutrient-dependent increase in pupal volume was smaller compared with female larvae (Fig. 1A; sex:diet interaction *p* <0.0001; S1 Table). This suggests that in nutrient-rich conditions, there is a sex difference in body size plasticity, where nutrient-dependent phenotypic plasticity is higher in females. To represent the normal body size responses of each sex to nutrient quantity, we plotted reaction norms for pupal volume in *w^1118^* males and females raised on different diets (Fig. 1B). The body size response to increased nutrient quantity between 0.5× and 1× was not different between the sexes (Fig. 1B); however, the body size response to increased nutrient quantity between 1× and 2× was larger in females than in males (Fig. 1B). Importantly, these findings were not specific to pupal volume, as we reproduced our findings using adult weight as an additional readout for body size (Fig. 1C, D). Thus, our findings demonstrate that while phenotypic plasticity is similar between the sexes in some nutritional contexts, body size plasticity is higher in females than in males in a nutrient-rich environment.

**Figure 1.**
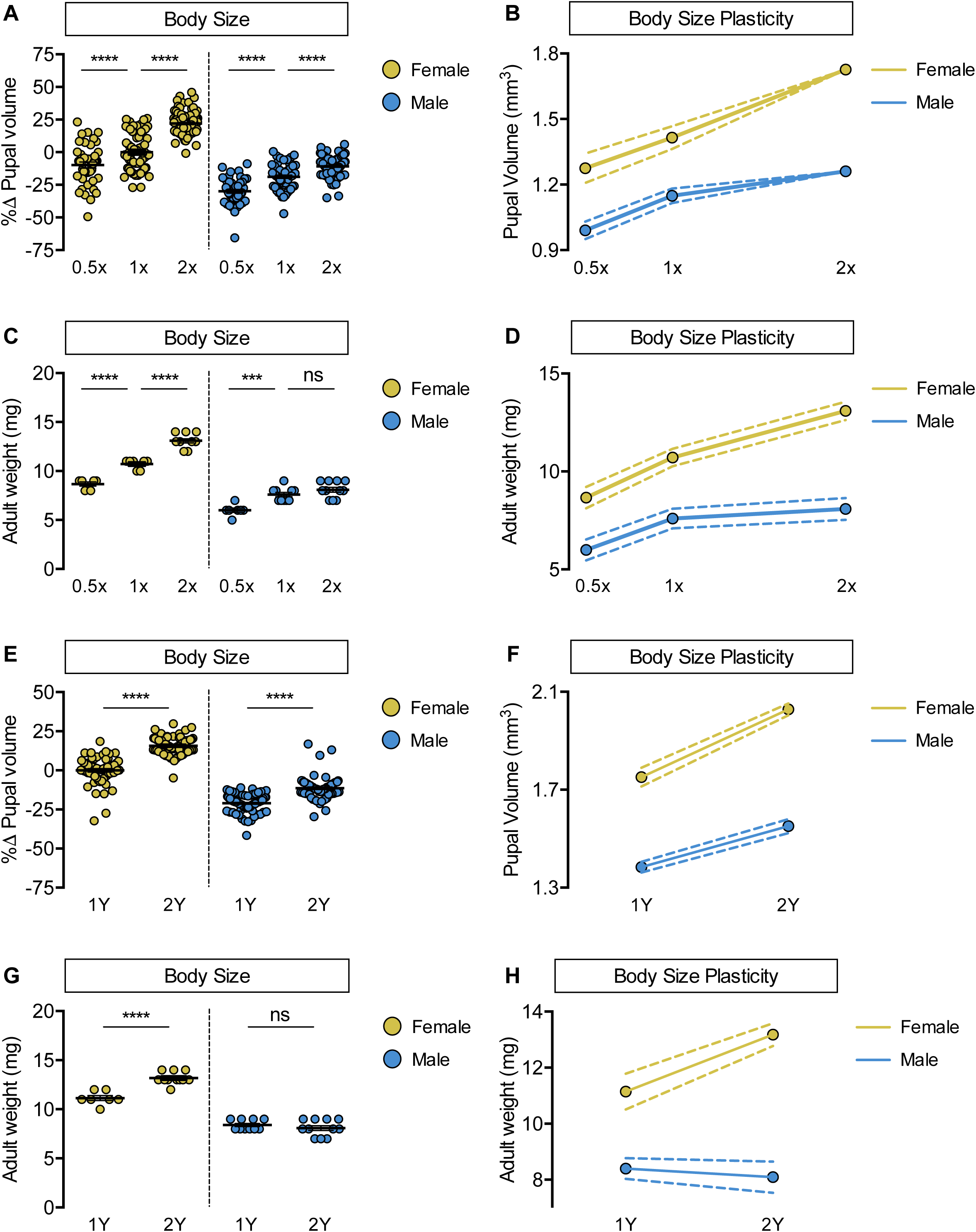
Increased female body size plasticity in a protein-rich diet. (A) Pupal volume was significantly higher in *w^1118^* males and females cultured on a widely-used diet (1×) compared with larvae raised on a reduced-nutrient diet (0.5×) (*p*<0.0001 and *p* = 0.0006, respectively; two-way ANOVA followed by Tukey HSD test). The magnitude of this increase in pupal volume was the same in both sexes (sex:diet interaction *p* = 0.7048; two-way ANOVA followed by Tukey HSD test). Pupal volume was significantly higher in *w^1118^* males and females raised on a nutrient-rich diet (2×) compared with larvae cultured on 1× (*p*<0.0001 for both; two-way ANOVA followed by Tukey HSD test); however, the magnitude of the increase in body size was significantly larger in females than in males (sex:diet interaction *p*<0.0001; two-way ANOVA followed by Tukey HSD test). (B) Reaction norms for pupal volume in *w^1118^* larvae raised on diets of varying quantity (0.5×, 1×, 2×), plotted using data presented in panel A. n = 43-100 pupae. (C) Adult weight was significantly higher in *w^1118^* males and females cultured on 1× compared with flies raised on 0.5× (*p*<0.0001 for both sexes; two-way ANOVA followed by Tukey HSD test). The magnitude of this increase in adult weight was the same in both sexes (sex:diet interaction *p* = 0.3197; two-way ANOVA followed by Tukey HSD test). Adult weight was significantly higher in *w^1118^* females raised on 2× compared to flies cultured on 1×; however, male adult weight was not significantly increased (*p*<0.0001 and *p* = 0.4015, respectively; two-way ANOVA followed by Tukey HSD test), where the diet-dependent increase in adult weight was higher in females (sex:diet interaction *p* = 0.0003; two-way ANOVA followed by Tukey HSD test). (D) Reaction norms for adult weight in response to changes in nutrient quantity in *w^1118^* females and males, plotted using the data presented in panel C. n = 6-11 groups of 10 flies. (E) Pupal volume was significantly higher in both males and females cultured on a yeast-rich medium (2Y) compared with larvae raised on a diet containing half the quantity of yeast (1Y) (*p*<0.0001 for both sexes; two-way ANOVA followed by Tukey HSD test); however, the magnitude of the nutrient-dependent increase in pupal volume was larger in females than in males (sex:diet interaction *p* = 0.0001; two-way ANOVA followed by Tukey HSD test). (F) Reaction norms for pupal volume in response to changes in dietary yeast in *w^1118^* females and males, plotted using the data in panel E. n = 62-80 pupae. (G) Adult weight was significantly higher in females cultured on 2Y compared with flies raised on 1Y; however, male adult weight was not significantly higher in flies raised on 2Y compared with males cultured on 1Y (*p*<0.0001 and *p* = 0.7199, respectively; two-way ANOVA followed by Tukey HSD test, sex:diet interaction *p*<0.0001). (H) Reaction norms for adult weight in *w^1118^* females and males reared on either 1Y or 2Y, plotted using data from panel G. n = 7-11 groups of 10 flies. For body size plasticity graphs, filled circles indicate mean body size, and dashed lines indicate 95% confidence interval. *** indicates p<0.001, **** indicates p<0.0001; ns indicates not significant; error bars indicate SEM.

To narrow down macronutrients that account for the increased body size plasticity in females, we changed individual food ingredients and measured body size in *w^1118^* males and females. We first altered dietary yeast, as previous studies show that yeast is a key source of protein and an important determinant of larval growth (Britton et al., 2002; Géminard et al., 2009; Robertson, 1963). In *w^1118^* females raised on a diet with yeast content that corresponds to the amount in the 2× diet (2Y diet), pupal volume was significantly larger than in females raised on a diet containing half the yeast content (1Y) (Fig. 1E). It is important to note that the yeast content of the 1Y diet is within the range of many larval growth studies (Ghosh et al., 2014; Koyama & Mirth, 2016; Marshall et al., 2012; Sawala & Gould, 2017), and therefore does not represent a nutrient-restricted diet. In *w^1118^* males, the magnitude of the nutrient-dependent increase in pupal volume was smaller than in females (Fig. 1E; sex:diet interaction *p* = 0.0001; S1 Table), suggesting that nutrient-dependent body size plasticity is higher in females in a yeast-rich context. Indeed, when we plot reaction norms for pupal volume in both sexes, the magnitude of the yeast-dependent change in pupal volume (Fig. 1F) and adult weight (Fig. 1G, H) was larger in females than in males. This sex difference in phenotypic plasticity in a yeast-rich context was reproduced in *Canton-S* (*CS*), a wild-type strain (Fig. S1A, B), and using wing length as an additional measure of growth (Fig. S2A). Thus, our findings indicate that the male-female difference in nutrient-dependent body size plasticity persists across multiple genetic backgrounds, and confirms that body size is a robust trait to monitor nutrient-dependent phenotypic plasticity.

Given the sex difference in body size plasticity in response to altered yeast content, we hypothesized that yeast may trigger increased nutrient-dependent body size plasticity in females. To test this, we raised larvae on diets with altered sugar (Fig. S3A) or calorie content (Fig. S3B). Because we observed no sex:diet interaction for either manipulation (sex:diet interaction *p* = 0.6536 and *p* = 0.3698, respectively; S1 Table), this suggests dietary yeast mediates the sex difference in nutrient-dependent body size plasticity. To test whether protein is the macronutrient in yeast that enables sex-specific phenotypic plasticity, we pharmacologically limited protein breakdown by culturing larvae on the 2Y diet supplemented with either a broad-spectrum protease inhibitor (protease inhibitor cocktail; PIC) or a serine protease-specific inhibitor (4-(2-aminoethyl)benzenesulfonyl fluoride hydrochloride; AEBSF). We found a significant body size reduction in both sexes treated with protease inhibitors (Fig. S4A, B), in line with previous studies (Erkosar et al., 2015); however, the inhibitor-induced decrease in pupal volume was larger in female larvae than in males (sex:treatment interaction *p* = 0.0029 [PIC] and *p*<0.0001 [AEBSF]; S1 Table). This indicates that yeast-derived dietary protein is the macronutrient that augments nutrient-dependent body size plasticity in females. While one potential explanation for the male-female difference in body size plasticity is a sex difference in food intake or length of the growth period, we found no male-female differences in either phenotype between *w^1118^* male and female larvae cultured on 1Y or 2Y (Fig. S5A-C). Moreover, the larger body size of female larvae does not explain their increased nutrient-dependent body size plasticity, as a genetic manipulation that augments male body size did not enhance phenotypic plasticity (Fig. S6A, B). Taken together, our data reveals female larvae have enhanced body size plasticity in a nutrient-rich context, and identifies abundant dietary protein as a prerequisite for females to maximize body size.

### A nutrient-dependent upregulation of IIS activity is required for body size plasticity in females

In a mixed-sex population of *Drosophila* larvae, IIS activity is positively regulated by nutrient availability to promote growth (Böhni et al., 1999; Britton et al., 2002; Chen et al., 1996; Fernandez et al., 1995; Grewal, 2009; Teleman, 2009). We therefore examined nutrient-dependent changes to IIS activity in larvae raised on 1Y and 2Y (Fig. 2A-D). Previous studies have shown that high levels of IIS activity repress mRNA levels of several genes, including *InR*, *brummer* (*bmm*, FBgn0036449), and *eukaryotic initiation factor 4E-binding protein* (*4E-BP*, FBgn0261560) (Alic et al., 2011; Jünger et al., 2003; Kang et al., 2017; Puig & Tjian, 2005; Zinke et al., 2002). In *w^1118^* females, we found that the mRNA levels of *InR*, *bmm*, and *4E-BP* were significantly lower in larvae reared on 2Y than in larvae raised on 1Y (Fig. 2A). This suggests IIS activity is significantly higher in females raised on 2Y than in females cultured on 1Y. To confirm this, we used the localization of a ubiquitously-expressed green fluorescent protein (GFP) fused to a pleckstrin homology (PH) domain (GFP-PH) as an additional readout of IIS activity. Because high levels of IIS activity raise the level of PIP_3_ at the plasma membrane, and PH domains bind specifically to PIP_3_, larvae with elevated IIS activity show increased membrane localization of GFP-PH (Britton et al., 2002). We observed a significantly higher membrane localization of GFP-PH in females cultured on 2Y than in female larvae raised on 1Y (Fig. 2B), indicating enhanced IIS activity in females raised on 2Y. In *w^1118^* males, we observed no significant difference in the mRNA levels of *InR*, *bmm,* and *4E-BP* between larvae grown on 2Y compared with larvae cultured on 1Y (Fig. 2C). Further, there was no significant difference in GFP-PH membrane localization between males raised on 2Y and males reared on 1Y (Fig. 2D). Together, these results suggest that IIS activity was enhanced by a protein-rich diet in female larvae but not in males, revealing a previously unrecognized sex difference in diet-induced changes to IIS activity.

**Figure 2.**
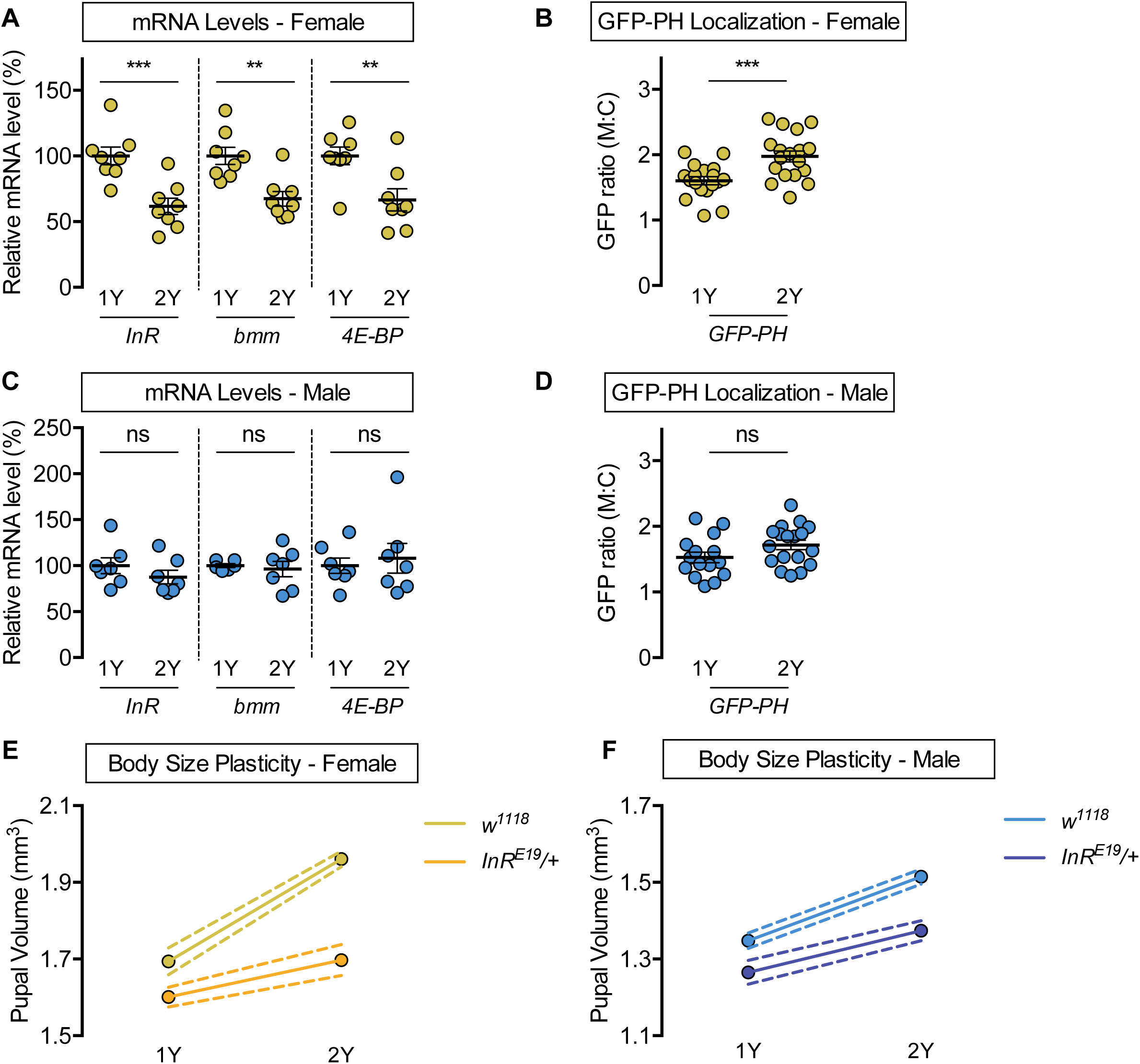
Upregulation of IIS activity is required for increased nutrient-dependent body size plasticity in females. (A) In females, mRNA levels of the *insulin receptor* (*InR*)*, brummer* (*bmm*), and *eukaryotic initiation factor 4E-binding protein* (*4E-BP*) were significantly lower in larvae raised on a protein-rich diet (2Y) compared with larvae raised on a diet containing half the protein content (1Y) (*p* = 0.0009, 0.0019, and 0.0077, respectively; Student’s *t* test). n = 8 biological replicates. (B) Quantification of the ratio between cell surface membrane-associated green fluorescent protein (GFP) and cytoplasmic GFP fluorescence (GFP ratio [M:C]) in a dissected fat body of female larvae from the GFP-PH strain. The GFP ratio was significantly higher in female larvae cultured on 2Y compared with larvae raised on 1Y (*p* = 0.001; Student’s *t* test). n = 18 biological replicates. (C) In males, there was no significant difference in *InR*, *bmm,* or *4E-BP* mRNA levels between larvae raised on 2Y compared with larvae cultured on 1Y (*p* = 0.291, 0.6994, and 0.666, respectively; Student’s *t* test). n = 6-7 biological replicates. (D) In males, the GFP ratio (M:C) was not significantly different between males cultured on 2Y compared with larvae raised on 1Y (*p* = 0.0892; Student’s *t* test). n = 15-18 biological replicates. (E) Pupal volume was significantly higher in both *w^1118^* females and *InR^E19^/+* females reared on 2Y compared with genotype-matched females cultured on 1Y (*p*<0.0001 for both genotypes; two-way ANOVA followed by Tukey HSD test); however, the magnitude of the nutrient-dependent increase in pupal volume was lower in *InR^E19^/+* females (genotype:diet interaction *p*<0.0001; two-way ANOVA followed by Tukey HSD test). n = 58-77 pupae. (F) Pupal volume was significantly higher in both *w^1118^* males and *InR^E19^/+* males reared on 2Y compared with genotype-matched males cultured on 1Y (*p*<0.0001 for both genotypes; two-way ANOVA followed by Tukey HSD test). While we observed a sex:diet interaction in the *w^1118^* control genotype, there was no sex:diet interaction in the *InR^E19^/+* genotype (*p*<0.0001 and *p* = 0.7104, respectively; two-way ANOVA followed by Tukey HSD test). n = 47-76 pupae. For body size plasticity graphs, filled circles indicate mean body size, and dashed lines indicate 95% confidence interval. ** indicates *p*<0.01, *** indicates *p*<0.001, ns indicates not significant; error bars indicate SEM.

To determine whether this increased IIS activity is required in females for the ability to maximize body size in response to dietary protein, we measured pupal volume in larvae heterozygous for a hypomorphic mutation in the *InR* gene (*InR^E19^/+*) that were raised in either 1Y or 2Y. Previous studies have shown that while overall growth is largely normal in *InR^E19^/+* heterozygous animals, growth that requires high levels of IIS activity is blunted (Chen et al., 1996; Rideout et al., 2012, 2015). In *w^1118^* control females, larvae cultured on 2Y were significantly larger than larvae raised on 1Y (Fig. 2E); however, the magnitude of this protein-dependent increase in pupal volume was smaller in *InR^E19^/+* females (Fig. 2E; genotype:diet interaction *p*<0.0001; S1 Table). This suggests that nutrient-dependent body size plasticity was reduced in *InR^E19^/+* females. Indeed, while we observed a sex difference in phenotypic plasticity in the *w^1118^* control genotype (sex:diet interaction *p*<0.0001 S1 Table), the sex difference in nutrient-dependent body size plasticity was abolished in the *InR^E19^/+* genotype (Fig. 2E, F: sex:diet interaction *p* = 0.7104; S1 Table). Together, these results indicate that the nutrient-dependent upregulation of IIS activity in females is required for their increased phenotypic plasticity, and suggest that the sex difference in body size plasticity arises from the female-specific ability to enhance IIS activity in a protein-rich context.

### *dilp2* is required for the nutrient-dependent upregulation of IIS activity and body size plasticity in females

Previous studies have identified changes to the production and release of dILPs as important mechanisms underlying nutrient-dependent changes to IIS activity and body size (Colombani et al., 2003; Géminard et al., 2009; Zhang et al., 2009). For example, the mRNA levels of *dilp3* and *dilp5*, but not *dilp2*, decrease in response to nutrient withdrawal (Colombani et al., 2003; Géminard et al., 2009; Ikeya et al., 2002), and the release of dILPs 2, 3, and 5 from the IPCs is altered by changes in nutrient availability (Géminard et al., 2009; Kim & Neufeld, 2015). Interestingly, a recent study showed that late third-instar female larvae have increased dILP2 secretion compared with age-matched males when the larvae were raised in a diet equivalent to 2Y (Rideout et al., 2015). Given that dILP2 is an important growth-promoting dILP (Grönke et al., 2010; Ikeya et al., 2002), we tested whether *dilp2* was required in females for the nutrient-dependent upregulation of IIS activity. In control *w^1118^* females, mRNA levels of *4E-BP* and *InR* were significantly lower in larvae raised on 2Y than in larvae reared on 1Y (Fig. 3A and Fig. S7A), suggesting a nutrient-dependent increase in IIS activity. In contrast, mRNA levels of *4E-BP* and *InR* were not significantly lower in *dilp2* female larvae raised on 2Y compared with genotype-matched females cultured on 1Y (Fig. 3A and Fig. S7A). In *w^1118^* and *dilp2* males, mRNA levels of *4E-BP* were not significantly lower in larvae raised on 2Y compared with genotype-matched larvae cultured on 1Y (Fig. 3B), trends we also observed using *InR* (Fig. S7B). This data suggests that *dilp2* is required for the nutrient-dependent upregulation of IIS activity in females in a protein-rich context.

**Figure 3.**
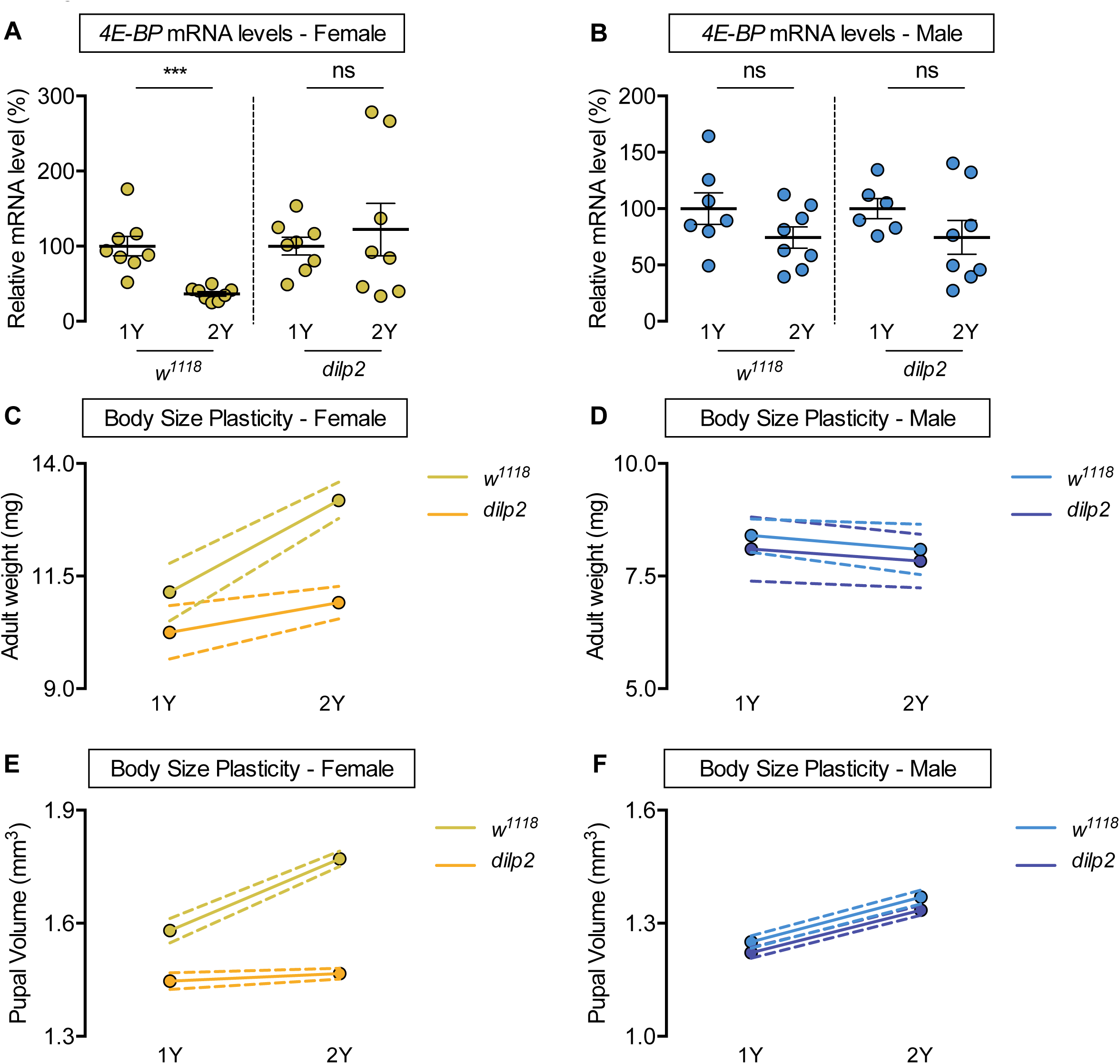
*Drosophila* insulin-like peptide 2 is required for the nutrient-dependent upregulation of insulin pathway activity and increased female body size plasticity. (A) In control *w^1118^* females, mRNA levels of *eukaryotic initiation factor 4E-binding protein (4E-BP)* were significantly lower in larvae cultured on a protein-rich diet (2Y) compared with larvae raised on a diet containing half the protein content (1Y) (*p* = 0.0003; Student’s *t* test). In *dilp2* mutant females, there was no significant difference in *4E-BP* mRNA levels in larvae cultured on 2Y compared with larvae raised on 1Y (*p* = 0.5563; Student’s *t* test). n = 8 biological replicates. (B) In control *w^1118^* and *dilp2* mutant males, mRNA levels of *4E-BP* were not significantly lower in larvae cultured on 2Y compared with larvae raised on 1Y (*p* = 0.1429 and *p* = 0.207, respectively; Student’s *t* test). n = 7-8 biological replicates. (C) Adult weight was significantly higher in *w^1118^* females raised on 2Y compared with flies cultured on 1Y (*p*<0.0001; two-way ANOVA followed by Tukey HSD test); however, adult weight was not significantly different between *dilp2* mutant females reared on 2Y versus 1Y (*p* = 0.1263; two-way ANOVA followed by Tukey HSD test). n = 7-11 groups of 10 flies. (D) Adult weight in control *w^1118^* and *dilp2* mutant males was not significantly higher in flies reared on 2Y compared with males raised on 1Y (*p* = 0.8366 and *p* = 0.8817, respectively; two-way ANOVA followed by Tukey HSD test). There was a significant sex:diet interaction in the control *w^1118^* genotype (*p*<0.0001), but not in the *dilp2* mutant genotype (*p* = 0.0827; two-way ANOVA followed by Tukey HSD test). n = 10-12 groups of 10 flies. (E) Pupal volume was significantly higher in *w^1118^* females but not in *dilp2* mutant females reared on 2Y compared with genotype-matched females cultured on 1Y (*p*<0.0001 and *p* = 0.6486 respectively; two-way ANOVA followed by Tukey HSD test). The magnitude of the nutrient-dependent increase in pupal volume was higher in *w^1118^* females (genotype:diet interaction *p*<0.0001; two-way ANOVA followed by Tukey HSD test). n = 74-171 pupae. (F) Pupal volume was significantly higher in *w^1118^* males and *dilp2* mutant males reared on 2Y compared with genotype-matched males cultured on 1Y (*p*<0.0001 for both genotypes; two-way ANOVA followed by Tukey HSD test). The magnitude of the nutrient-dependent increase in pupal volume was not different between genotypes (genotype:diet interaction *p* = 0.6891; two-way ANOVA followed by Tukey HSD test). n = 110-135 pupae. For all body size plasticity graphs, filled circles indicate mean body size, and dashed lines indicate 95% confidence interval. *** indicates *p*<0.001; ns indicates not significant; error bars indicate SEM.

To determine whether the inability to augment IIS activity on 2Y affects nutrient-dependent body size plasticity in females, we measured body size in *w^1118^* and *dilp2* mutant larvae cultured on either 1Y or 2Y. In *w^1118^* control females, adult weight was significantly higher in flies cultured on 2Y compared with flies raised on 1Y (Fig. 3C); however, this nutrient-dependent increase in adult weight was not observed in *dilp2* mutant females (Fig. 3C; genotype:diet interaction *p* = 0.0024; S1 Table). In *w^1118^* control males and *dilp2* mutant males, there was no significant increase in adult weight in flies raised on 2Y compared with genotype-matched flies cultured on 1Y (Fig. 3D; genotype:diet interaction *p* = 0.935; S1 Table). Indeed, in contrast to the sex difference in nutrient-dependent body size plasticity in the *w^1118^* genotype (sex:diet interaction *p*<0.0001; S1 Table), the sex difference in phenotypic plasticity was abolished in the *dilp2* mutant genotype (sex:diet interaction *p* = 0.0827; S1 Table). Importantly, we replicated all these findings using pupal volume (Fig. 3E, F), and reproduced the female-specific effects of *dilp2* loss by globally overexpressing a *UAS-dilp2-RNAi* transgene (Fig. S7C). Further, changes to *dilp* mRNA levels in males and females lacking *dilp2* (Fig. S8A, B), and protein-dependent changes to *dilp* mRNA levels (Fig. S9A, B), were similar in both sexes. Thus, our data reveals a previously unrecognized female-specific requirement for *dilp2* in triggering a nutrient-dependent increase in IIS activity and body size in a protein-rich context.

### A nutrient-dependent increase in *stunted* mRNA levels is required for enhanced IIS activity and body size plasticity in females

Nutrient-dependent changes in dILP secretion from the IPCs, and consequently IIS activity, are mediated by humoral factors that are regulated by dietary nutrients (Britton & Edgar, 1998; Delanoue et al., 2016; Koyama & Mirth, 2016; Rajan & Perrimon, 2012; Rodenfels et al., 2014; Sano et al., 2015). For example, in a mixed-sex population of larvae, dietary protein augments mRNA levels of *Growth-blocking peptides 1* and *2* (*Gbp1*, FBgn0034199; *Gbp2*, FBgn0034200), *CCHamide-2* (*CCHa2;* FBgn0038147), *unpaired 2* (*upd2;* FBgn0030904), and *sun* (Delanoue et al., 2016; Koyama & Mirth, 2016; Rajan & Perrimon, 2012; Sano et al., 2015). Increased levels of these humoral factors promote the secretion of IPC-produced dILPs to enhance IIS activity and growth (Delanoue et al., 2016; Koyama & Mirth, 2016; Meschi et al., 2019; Rajan & Perrimon, 2012; Sano et al., 2015). To determine whether any humoral factors contribute to the sex-specific increase in IIS activity in a protein-rich diet, we examined mRNA levels of each factor in larvae of both sexes raised on either 1Y or 2Y. In *w^1118^* females, *sun* mRNA levels in larvae reared on 2Y were significantly higher than in larvae cultured on 1Y (Fig. 4A). In contrast, mRNA levels of *Gbp1*, *Gbp2*, *CCHa2*, and *upd2* were not significantly higher in female larvae reared on 2Y compared with 1Y (Fig. S10A). Thus, while previous studies have shown that mRNA levels of all humoral factors were severely reduced by a nutrient-restricted diet or nutrient withdrawal (Delanoue et al., 2016; Koyama & Mirth, 2016; Rajan & Perrimon, 2012; Sano et al., 2015), our study suggests that for most factors, augmenting dietary protein beyond a widely-used level does not further enhance mRNA levels. In males, there was no significant increase in *sun* mRNA levels (Fig. 4B), or any other humoral factors (Fig. S10B), in larvae reared on 2Y compared with 1Y. Thus, there is a previously unrecognized sex difference in the regulation of *sun* mRNA levels in a protein-rich context. Given that *sun* has previously been shown to promote IIS activity by enhancing dILP2 secretion (Delanoue et al., 2016), we hypothesized that the female-specific increase in *sun* mRNA levels in 2Y triggers the nutrient-dependent upregulation of IIS activity in females. To test this, we overexpressed UAS-*sun-RNAi* in the larval fat body using *r4-GAL4*, and cultured the animals on either 1Y or 2Y. Importantly, overexpression of the *UAS-sun-RNAi* transgene significantly decreased *sun* mRNA levels in both sexes (Fig. S10C, D), where GAL4 expression was similar between the sexes in 1Y and 2Y (Fig. S10E). In control *r4>+* and *+>UAS-sun-RNAi* females, we observed a significant decrease in *InR, bmm,* and *4E-BP* mRNA levels in larvae cultured on 2Y compared with genotype-matched larvae reared on 1Y (Fig. 4C). In contrast, the nutrient-dependent decrease in *InR, bmm,* and *4E-BP* mRNA levels was absent in *r4>UAS-sun-RNAi* females (Fig. 4C). In *r4>+*, *+>UAS-sun-RNAi*, and *r4>UAS-sun-RNAi* males, we found no consistent indications of increased IIS activity in larvae cultured on 2Y compared with genotype-matched larvae raised on 1Y (Fig. S11A). Together, this data suggests that in females a protein-rich diet stimulates a nutrient-dependent increase in *sun* mRNA that enhances IIS activity. In males, the 2Y diet did not augment *sun* mRNA levels, suggesting one reason for the lack of a nutrient-dependent increase in IIS activity.

**Figure 4.**
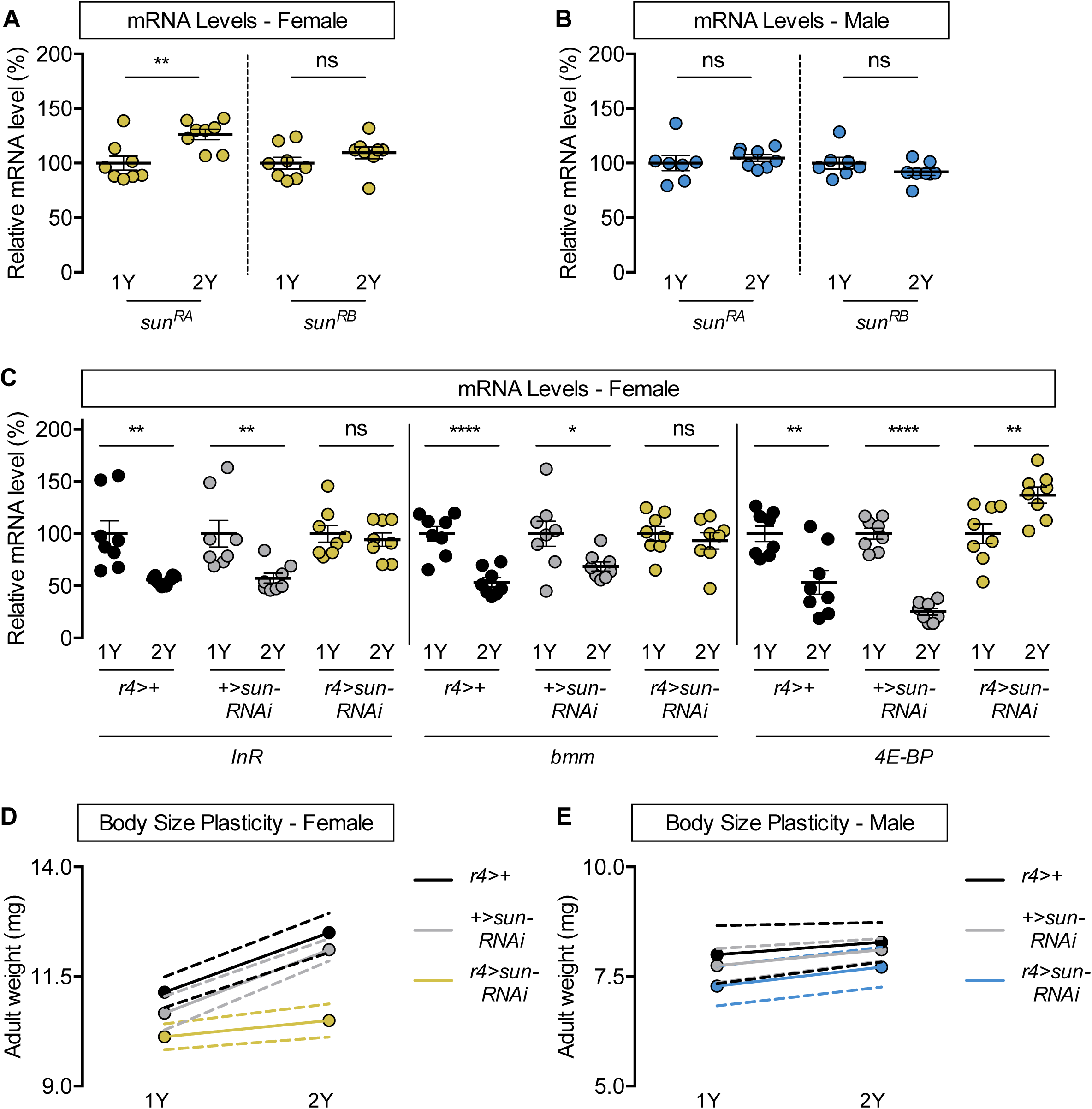
*stunted* is required for the nutrient-dependent upregulation of insulin pathway activity and increased female body size plasticity. (A) In females, mRNA levels of *stunted* (*sun*)*^RA^*, but not *sun^RB^,* were significantly higher in larvae cultured on a protein-rich diet (2Y) compared with larvae raised on a diet containing half the protein content (1Y) (*p* = 0.0055 and *p* = 0.2327, respectively; Student’s *t* test). n = 8 biological replicates. (B) In males, mRNA levels of *sun^RA^* and *sun^RB^* were not significantly different in larvae raised on 2Y compared with larvae raised on 1Y (*p* = 0.5832 and *p* = 0.2017, respectively; Student’s *t* test). n = 7-8 biological replicates. (C) In control *r4>+* and *+>UAS-sun-RNAi* females, mRNA levels of the *insulin receptor* (*InR*)*, brummer* (*bmm*), and *eukaryotic initiation factor 4E-binding protein* (*4E-BP*) mRNA levels were significantly lower in larvae cultured on 2Y compared with larvae raised on 1Y (*p* = 0.0032, *p*<0.0001, and 0.0041 [*r4>+*], and 0.0074, 0.0281, *p*<0.0001 [*+>UAS-sun-RNAi*], respectively; Student’s *t* test). In contrast, mRNA levels of *InR* and *bmm* were not significantly different in *r4>UAS-sun-RNAi* females raised on 2Y compared with genotype-matched females reared on 1Y (*p* = 0.5897 and *p* = 0.5297, respectively; Student’s *t* test) and levels of *4E-BP* were significantly higher (*p* = 0.0094; Student’s *t* test). n = 8 biological replicates. (D) Adult weight was significantly higher in female flies raised in 2Y compared with females raised in 1Y in *r4>+* and *+>UAS-sun-RNAi* controls (*p*<0.0001 for both genotypes; two-way ANOVA followed by Tukey HSD test); however, adult weight was not significantly different between *r4>UAS-sun-RNAi* females reared on 2Y compared with genotype-matched females raised on 1Y (*p* = 0.5035; two-way ANOVA followed by Tukey HSD test). n = 7-10 groups of 10 flies. (E) Adult weight was not significantly higher in male flies reared in 2Y compared with males cultured in 1Y for *r4>+* and *+>UAS-sun-RNAi* controls or *r4>UAS-sun-RNAi* males (*p* = 0.8883, 0.6317, and 0.554, respectively; two-way ANOVA followed by Tukey HSD test). There was a significant sex:diet interaction in the *r4>+* and *+>UAS-sun-RNAi* control genotypes (*p* = 0.011 and *p* = 0.0005, respectively; two-way ANOVA followed by Tukey HSD test), but no sex:diet interaction in the *r4>UAS-sun-RNAi* genotype (*p* = 0.8749; two-way ANOVA followed by Tukey HSD test). n = 6-9 groups of 10 flies. For all body size plasticity graphs, filled circles indicate mean body size, and dashed lines indicate 95% confidence interval. * indicates *p*<0.05, ** indicates *p*<0.01, **** indicates *p*<0.0001; ns indicates not significant; error bars indicate SEM.

We next asked whether the female-specific increase in *sun* and its regulation of IIS activity contribute to nutrient-dependent body size plasticity. In *r4>+* and *+>UAS-sun-RNAi* control females, adult weight was significantly higher in flies cultured on 2Y compared with genotype-matched flies raised on 1Y (Fig. 4D). In contrast, the nutrient-dependent increase in adult weight was abolished in *r4>UAS-sun-RNAi* females (Fig. 4D; genotype:diet interaction *p* = 0.0014; S1 Table). This indicates *r4>UAS-sun-RNAi* females have reduced nutrient-dependent body size plasticity, a finding we confirmed using pupal volume (Fig. S11B). In *r4>+*, *+>UAS-sun-RNAi*, and *r4>UAS-sun-RNAi* male flies raised on 2Y, adult weight was not significantly higher than in genotype-matched males raised on 1Y (Fig. 4E; genotype:diet interaction *p* = 0.9278; S1 Table). Additionally, we replicated all these findings using pupal volume (Fig. S11C). Importantly, in contrast to the sex difference in nutrient-dependent body size plasticity we observed in the *r4>+* and *+>UAS-sun-RNAi* control genotypes (sex:diet interaction *p* = 0.011 and *p =* 0.0005, respectively; S1 Table), the sex difference in phenotypic plasticity was abolished in the *r4>UAS-sun-RNAi* genotype (sex:diet interaction *p* = 0.8749; S1 Table). This suggests that the female-specific increase in *sun* mRNA levels is required for the sex difference in nutrient-dependent plasticity. A sex-specific role for *sun* was further supported by the fact that we reproduced the female-specific effects of *sun* knockdown on body size using an additional GAL4 line (Fig. S12A), and by the fact that no other humoral factors caused sex-specific effects on body size (Fig. S12B, C). Further, while we show that fat body-specific *sun* overexpression was sufficient to increase body size in both sexes (Fig. S13A, B), body size plasticity in these larger males was not significantly different from control males (genotype:diet interaction *p* = 0.4959, S1 Table), in line with our earlier data showing that augmenting body size in males was not sufficient to confer phenotypic plasticity (Fig. S6B). Thus, our data suggests that the female-specific ability to upregulate *sun* in the 2Y diet enhances IIS activity to promote a larger body size, revealing the mechanism by which females, and not males, augment body size in a protein-rich context.

### Sex determination gene *transformer* promotes nutrient-dependent body size plasticity in females

We next investigated the increased ability of females to enhance IIS activity and augment body size in a protein-rich context. Given that previous studies have implicated sex determination gene *tra* in regulating body size in a diet equivalent to the 2Y diet (Rideout et al., 2015), and identified links between *tra* and IIS activity in this context (Rideout et al., 2015), we explored a role for *tra* in regulating the sex difference in the nutrient-dependent upregulation of IIS activity and body size plasticity. In control *w^1118^* females, *4E-BP* mRNA levels were significantly lower in larvae raised on 2Y compared with larvae cultured on 1Y (Fig. 5A); however, this nutrient-dependent decrease in *4E-BP* mRNA levels was absent in *tra* mutant females (*tra^1^/Df(3L)st-j7)* (Fig. 5A). Similarly, while *sun* mRNA levels in *w^1118^* control females were significantly higher in larvae raised on 2Y compared with 1Y (Fig. 5B), this nutrient-dependent increase in *sun* mRNA levels was absent in *tra* mutant females (Fig. 5B). Thus, *tra* is required in females for the nutrient-dependent increase in *sun* mRNA and IIS activity in a protein-rich context. To determine whether the inability of *tra* mutant females to upregulate *sun* mRNA levels and IIS activity impacts nutrient-dependent body size plasticity, we measured body size in *w^1118^* controls and *tra* mutants raised in 1Y and 2Y. In control *w^1118^* females, adult weight was significantly higher in flies raised on 2Y compared with flies cultured on 1Y (Fig. 5C); however, this nutrient-dependent increase in adult weight was not observed in *tra* mutant females (Fig. 5C; genotype:diet interaction *p*<0.0001; S1 Table), a finding we reproduced using pupal volume (Fig. S14A). This indicates that *tra* mutant females have reduced nutrient-dependent body size plasticity compared with control females. In control *w^1118^* and *tra* mutant males, adult weight was not significantly higher in flies raised on 2Y compared with genotype-matched flies reared on 1Y (Fig. 5D; genotype:diet interaction *p* = 0.4507). Importantly, we replicated all these findings using pupal volume (Fig. S14B). Given that we observed a sex difference in nutrient-dependent body size plasticity in the *w^1118^* genotype (sex:diet interaction *p*<0.0001; S1 Table), but not in the *tra* mutant strain (sex:diet interaction *p* = 0.6598; S1 Table), our data reveals a previously unrecognized requirement for *tra* in regulating the sex difference in nutrient-dependent phenotypic plasticity.

**Figure 5.**
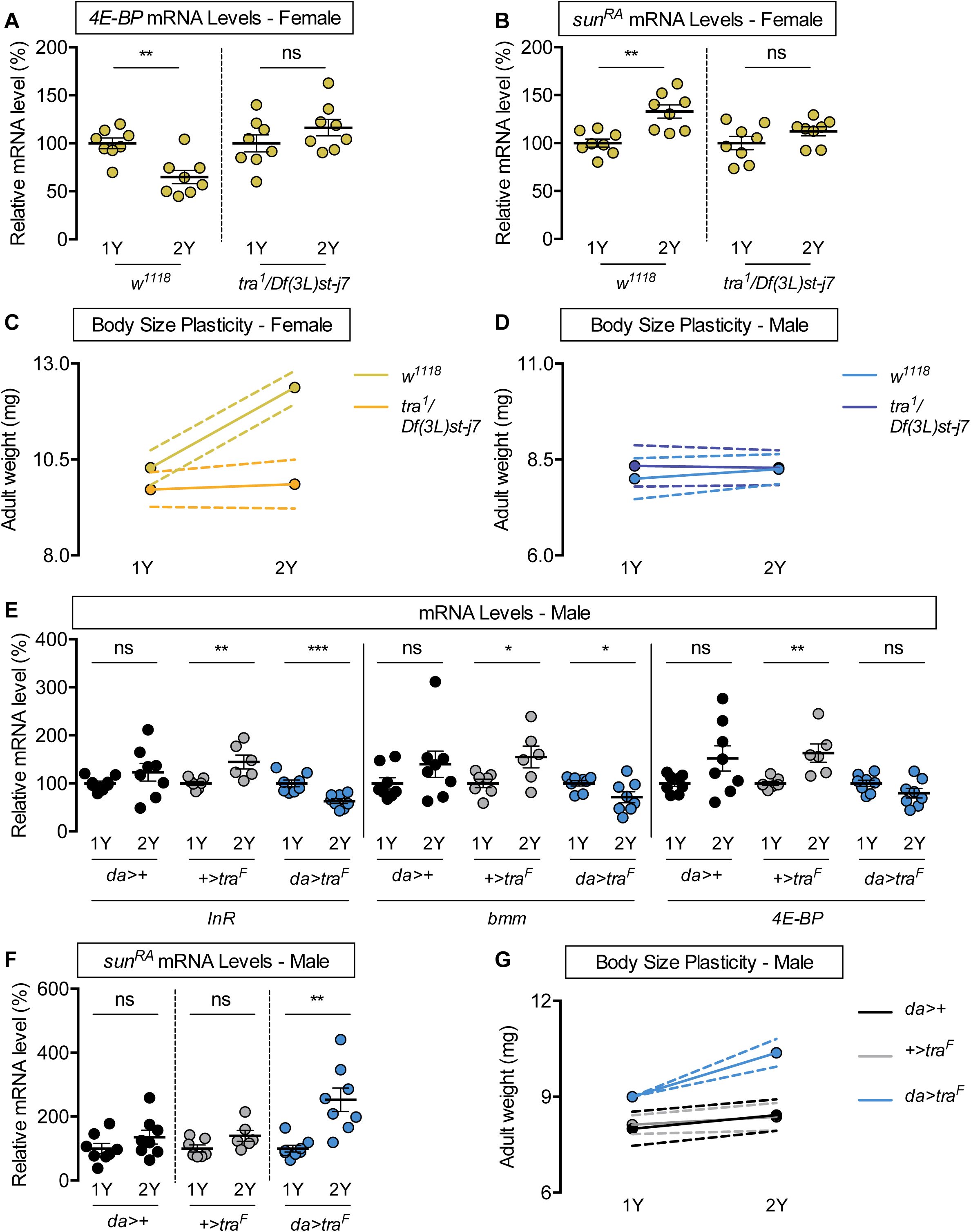
Sex determination gene *transformer* regulates increased nutrient-dependent body size plasticity in females. (A) In control *w^1118^* females, mRNA levels of *eukaryotic initiation factor 4E-binding protein (4E-BP)* were significantly lower in larvae cultured on a protein-rich diet (2Y) compared with larvae raised on a diet containing half the protein content (1Y) (*p* = 0.0013; Student’s *t* test). In *tra^1^*/*Df(3L)st-j7* females, there was no significant difference in *4E-BP* mRNA levels between larvae cultured on 2Y compared with larvae raised on 1Y (*p* = 0.2095; Student’s *t* test). n = 8 biological replicates (B) In control females, mRNA levels of *sun^RA^* were significantly higher in larvae cultured on 2Y compared with larvae raised on 1Y (*p* = 0.0011; Student’s *t* test); however, in *tra^1^*/*Df(3L)st-j7* females there was no significant difference in *sun^RA^* mRNA levels between larvae cultured on 2Y compared with larvae raised on 1Y (*p* = 0.1644; Student’s *t* test). n = 8 biological replicates. (C) Adult weight was significantly higher in *w^1118^* females raised on 2Y compared with females reared on 1Y (*p*<0.0001; two-way ANOVA followed by Tukey HSD test); however, there was no significant difference in adult weight between *tra^1^*/*Df(3L)st-j7* females cultured on 2Y compared with genotype-matched females raised on 1Y (*p* = 0.9617; two-way ANOVA followed by Tukey HSD test). n = 7-8 groups of 10 flies. (D) Adult weight was not significantly higher in either *w^1118^* control or *tra^1^*/*Df(3L)st-j7* mutant males in flies raised on 2Y compared with males reared on 1Y (*p* = 0.7808 and *p* = 0.9983, respectively; two-way ANOVA followed by Tukey HSD test). There was a significant sex:diet interaction in the *w^1118^* control genotype (*p*<0.0001; two-way ANOVA followed by Tukey HSD test); however, there was no sex:diet interaction in the *tra^1^*/*Df(3L)st-j7* genotype (*p* = 0.6598; two-way ANOVA followed by Tukey HSD test). n = 6-8 groups of 10 flies. (E) In control *da>+* males, mRNA levels of the *insulin receptor* (*InR*), *brummer* (*bmm*), and *4E-BP* were not significantly different between larvae cultured on 2Y compared with larvae raised on 1Y (*p* = 0.2418, 0.2033, and 0.0769, respectively; Student’s *t* test). In *+>UAS-tra^F^* males, mRNA levels of *InR, bmm,* and *4E-BP* were significantly increased between larvae cultured on 2Y compared with larvae raised on 1Y (*p* = 0.0088, 0.035, and 0.0052, respectively; Student’s *t* test). In *da>UAS-tra^F^* males, mRNA levels of *InR* and *bmm* were significantly lower in larvae cultured on 2Y compared with larvae raised on 1Y (*p* = 0.0007 and 0.0388, respectively; Student’s *t* test), and levels of *4E-BP* were not significantly altered (p = 0.103; Student’s *t* test). n = 6-8 biological replicates. (F) In control *da>+* and *+>UAS-tra^F^* males, mRNA levels of *sun^RA^* were not significantly different between larvae cultured on 2Y compared with larvae raised on 1Y (*p* = 0.2064 and *p* = 0.0711, respectively; Student’s *t* test). In contrast, *da>UAS-tra^F^* males showed a significant increase in mRNA levels of *sun^RA^* in larvae cultured on 2Y compared with males raised on 1Y (*p* = 0.0013; Student’s *t* test). n = 6-8 biological replicates. (G) Adult weight was not significantly higher in *da*>+ and *+>UAS-tra^F^* control males reared on 2Y compared with genotype-matched males flies cultured on 1Y (*p* = 0.5186 and *p* = 0.8858, respectively; two-way ANOVA followed by Tukey HSD test); however, there was a significant increase in adult weight between *da>UAS-tra^F^* males cultured on 2Y compared with genotype-matched flies raised on 1Y (*p*<0.0001; two-way ANOVA followed by Tukey HSD test). n = 7-8 groups of 10 flies. For body size plasticity graphs, filled circles indicate mean body size, and dashed lines indicate 95% confidence interval. * indicates *p*<0.05, ** indicates *p*<0.01, *** indicates *p*<0.001; ns indicates not significant; error bars indicate SEM.

To determine whether lack of a functional Tra protein in males explains their reduced nutrient-dependent body size plasticity, we overexpressed *UAS-tra^F^* in all tissues using *daughterless* (*da*)-*GAL4.* We first asked whether *tra* overexpression impacted the nutrient-dependent regulation of *sun* mRNA and IIS activity. In control *da>+* and *+>UAS-tra^F^* males, there was no significant decrease in *InR*, *bmm*, or *4E-BP* mRNA levels in larvae reared in 2Y compared with larvae raised in 1Y (Fig. 5E). In contrast, there was a significant nutrient-dependent decrease in mRNA levels of *InR* and *bmm* in *da>UAS-tra^F^* males (Fig. 5E). Similarly, while *sun* mRNA levels in control *da>+* and *+>UAS-tra^F^* males were not significantly higher in larvae raised on 2Y compared with larvae reared on 1Y (Fig. 5F), there was a nutrient-dependent increase in *sun* mRNA levels in *da>UAS-tra^F^* males (Fig. 5F). This suggests the presence of a functional Tra protein in males confers the ability to upregulate *sun* mRNA levels and IIS activity in a protein-rich context. Next, we asked whether expressing a functional Tra protein in males would augment nutrient-dependent body size plasticity. In control *da>+* and *+>UAS-tra^F^* males, there was no significant increase in adult weight in flies raised on 2Y compared with genotype-matched flies reared on 1Y (Fig. 5G); however, there was a nutrient-dependent increase in *da>UAS-tra^F^* males (Fig. 5G; genotype:diet interaction *p* = 0.0038; S1 Table), a finding we reproduced using pupal volume (Fig. S15A). Thus, *da>UAS-tra^F^* males have increased phenotypic plasticity compared with control males, revealing a new role for *tra* in regulating nutrient-dependent body size plasticity. In females, we observed a significant increase in both adult weight and pupal volume in *da>+*, *+>UAS-tra^F^*, and *da>UAS-tra^F^* flies raised on the 2Y diet compared with genotype-matched females cultured on the 1Y diet (Fig. S15B, C). Because one study suggested high levels of Tra overexpression could cause lethality (Siera & Cline, 2008), we reproduced these findings using a recently published strain of flies in which adult males and females lacking *tra* (*tra^KO^*), and adult males and females carrying a cDNA encoding the female-specific Tra protein knocked into the *tra* locus (*tra^F K-IN^*), are produced from the same cross (Hudry et al., 2016, 2019). In line with *tra^1^/Df(3L)st-j7* females, *tra^KO^* females had reduced body size plasticity compared with control *w^1118^* and *tra^F K-IN^* females in a protein-rich context (Fig. S15D; genotype:diet interaction *p*<0.0001 S1 Table). As with *da>UAS-tra^F^* males, we found that *tra^F K-IN^* males, which express physiological levels of a functional Tra protein, showed increased nutrient-dependent body size plasticity compared with control *w^1118^* and *tra^KO^* males (Fig. S15E; genotype:diet interaction *p*<0.0001; S1 Table). Importantly, the sex difference in nutrient-dependent body size plasticity that we observed in the *w^1118^* genotype (sex:diet interaction p<0.0001) was abolished in the *tra^KO^* and *tra^F K-IN^* genotypes (*p* = 0.5068 and *p* = 0.3168, respectively; S1 Table). Together, our findings reveal a new role for *tra* in regulating the sex difference in nutrient-dependent body size plasticity.

These findings suggest that a functional Tra protein confers the ability to adjust body size in a protein-rich context via regulation of *sun* mRNA and IIS activity. To test this, we examined whether the ability to adjust *sun* mRNA levels is required for Tra’s effects on phenotypic plasticity. Because animals homozygous for null mutations in *sun* are larval lethal (Kidd et al., 2005), and *sun* is located on the X chromosome which precludes studies on flies heterozygous for a *sun* mutant allele, we examined nutrient-dependent body size plasticity in *da>UAS-tra^F^* animals heterozygous for a hypomorphic allele of *spargel* (*srl*, FBgn0037248), the *Drosophila* homolog of *peroxisome proliferator-activated receptor gamma coactivator 1-alpha* (*PGC-1α*). A previous study showed that *srl/PGC-1α*, an essential gene, was required for normal *sun* mRNA levels (Delanoue et al., 2016). Therefore, we predicted that heterozygous loss of *srl/PGC-1α* would blunt the nutrient-dependent increase in *sun* mRNA levels without compromising viability. While adult weight in *da>UAS-tra^F^* males and females was significantly higher in flies raised on 2Y compared with flies cultured in 1Y (Fig. 6A, B), as in Fig. 5G and Fig. S15C, the nutrient-dependent increase in adult weight was abolished in *da>UAS-tra^F^* males and females carrying a mutant allele of *srl/PGC-1α* (*srl^1^*) (Fig. 6A, B; genotype:diet interaction *p* = 0.0146 and *p* = 0.0008, respectively). This finding suggests that nutrient-dependent body size plasticity was reduced in *da>UAS-tra^F^,srl^1^/+* flies compared with controls. Therefore, when taken together, our results indicate that the nutrient-dependent upregulation of *sun* is important for *tra*’s ability to promote growth in a protein-rich context, revealing one mechanism by which Tra regulates body size plasticity.

**Figure 6.**
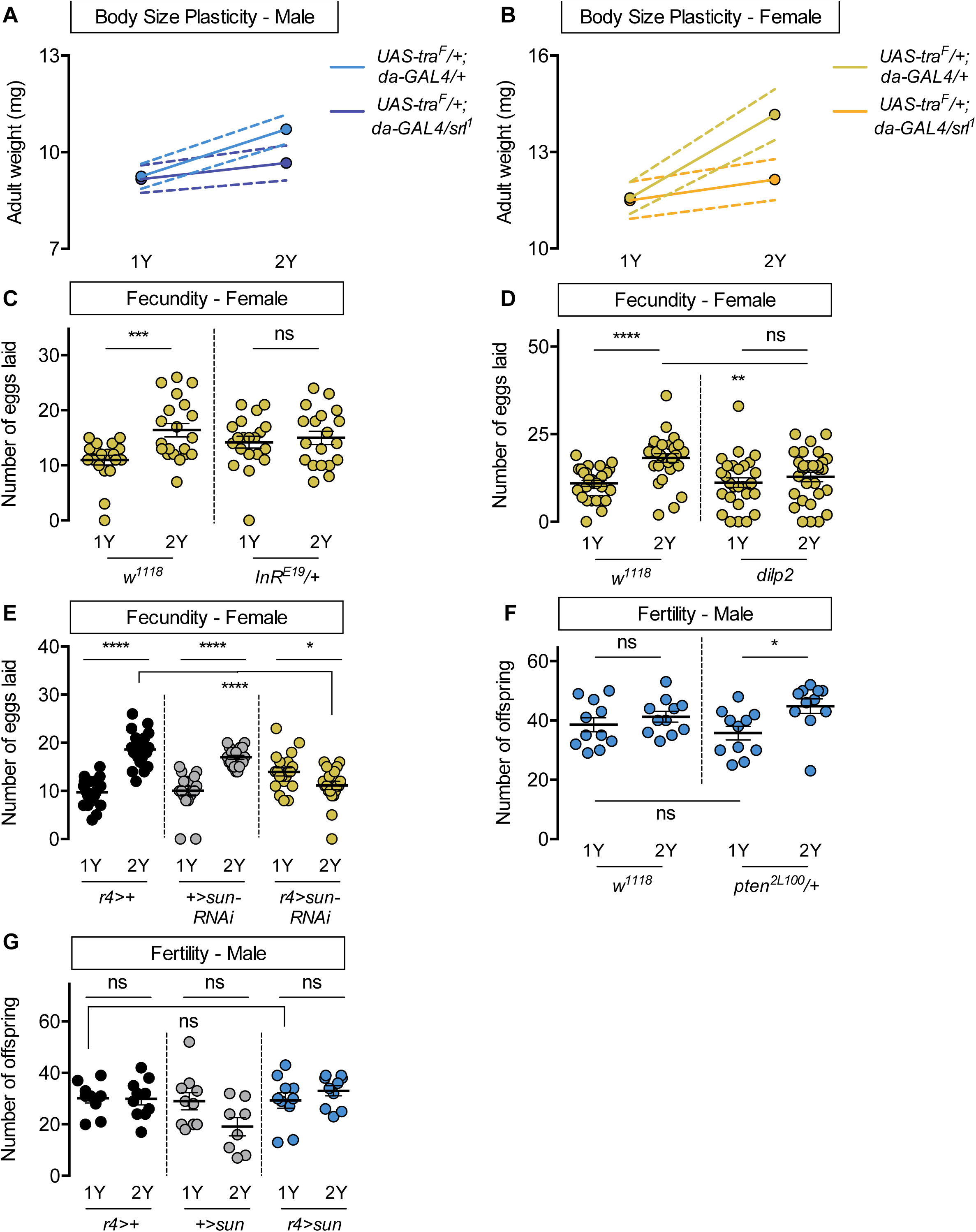
Increased nutrient-dependent body size plasticity in females promotes fertility. (A) Adult weight was higher in *da>UAS-tra^F^* males raised on a protein-rich diet (2Y) compared with *da>UAS-tra^F^* males reared on a diet containing half the protein content (1Y) (*p*<0.0001; two-way ANOVA followed by Tukey HSD test). In contrast, the nutrient-dependent increase in adult weight was abolished in *da>UAS-tra^F^* males heterozygous for a loss-of-function allele of *spargel (srl^1^)* (*p* = 0.2811; two-way ANOVA followed by Tukey HSD test). n = 6-8 groups of 10 flies. (B) Adult weight was higher in *da>UAS-tra^F^* females raised on 2Y compared with *da>UAS-tra^F^* females reared on 1Y (*p*<0.0001; two-way ANOVA followed by Tukey HSD test). In contrast, the nutrient-dependent increase in adult weight was absent in *da>UAS-tra^F^* females heterozygous for *srl^1^* (*p* = 0.2927; two-way ANOVA followed by Tukey HSD test). n = 6-7 groups of 10 flies. (C) In control *w^1118^* females there was a significant increase in the number of eggs laid by females raised on 2Y compared with females cultured on 1Y (*p* = 0.0009; Student’s *t* test); however, there was no significant difference in the number of eggs laid between *InR^E19^/+* females cultured on 2Y compared with genotype-matched females raised on 1Y (*p* = 0.617; Student’s *t* test). n = 19-20 biological replicates. (D) In control *w^1118^* females, there was a significant increase in the number of eggs laid by females raised on 2Y compared with females cultured on 1Y (*p*<0.0001; Student’s *t* test); however, there was no significant difference in the number of eggs laid between *dilp2* mutant females cultured on 2Y compared with females raised on 1Y (*p* = 0.4105; Student’s *t* test). n = 28-30 biological replicates. (E) In control *r4>+* and *+>UAS-sun-RNAi* females there was a significant increase in the number of eggs laid by females raised on 2Y compared with control females cultured on 1Y (*p*<0.0001 for both genotypes; Student’s *t* test). In *r4>UAS-sun-RNAi* females, the number of eggs laid by females cultured on 2Y was lower than females raised on 1Y (*p* = 0.0243; Student’s *t* test). n = 20 biological replicates. (F) In control *w^1118^* males there was no significant difference in the number of offspring produced between a 1Y and 2Y diet (*p* = 0.3662; Student’s *t* test). There was also no significant difference in the number of offspring produced between control *w^1118^* males and males heterozygous for a loss-of-function allele of *phosphatase and tensin homolog* (*pten;* genotype *pten^2L100^/+*) raised on 1Y (*p* = 0.4003; Student’s *t* test). Unlike control males, *pten^2L100^/+* males reared on 2Y produced significantly more offspring than genotype-matched males raised on 1Y (*p* = 0.0137; Student’s *t* test). n = 11 biological replicates. (G) In control *r4>+* and *+>UAS-sun* and experimental *r4>UAS-sun* males, there was no significant effect on the number of offspring produced between a 1Y and 2Y diet (*p* = 0.9222, 0.0595, and 0.32 respectively; Student’s *t* test). There was also no significant difference in the number of offspring produced between control *r4>+, +>UAS-sun* males and experimental *r4>UAS-sun* males raised on 1Y (*p* = 0.9723 and *p* = 0.9969 respectively; one-way ANOVA followed by Tukey HSD test). n = 8-10 groups of 10 flies. For body size plasticity graphs, filled circles indicate mean body size, and dashed lines indicate 95% confidence interval. * indicates *p*<0.05, *** indicates *p*<0.001, **** indicates *p*<0.0001; ns indicates not significant; error bars indicate SEM.

### Increased nutrient-dependent body size plasticity in females promotes fecundity in a protein-rich context

Previous studies have shown that plentiful nutrients during development maximize body size to promote fertility in *Drosophila* females (Bergland et al., 2008; Green & Extavour, 2014; Grönke et al., 2010; Hodin & Riddiford, 2000; Mendes & Mirth, 2016; Robertson, 1957a, 1957b; Sarikaya et al., 2012; Tu & Tatar, 2003), and that high levels of IIS activity are required for normal egg development, ovariole number, and fecundity (Green & Extavour, 2014; Grönke et al., 2010; Mendes & Mirth, 2016; Richard et al., 2005). In line with these findings, *w^1118^* female flies reared on 2Y produced significantly more eggs compared with genotype-matched females cultured on 1Y (Fig. 6C). This suggests that the ability to augment IIS activity and body size in response to a protein-rich diet allows females to maximize fecundity in conditions where nutrients are plentiful. To test this, we measured the number of eggs produced by *InR^E19^*/+ females and *w^1118^* controls raised in either 1Y or 2Y. In contrast to *w^1118^* females, the nutrient-dependent increase in egg production was absent in *InR^E19^*/+ females (Fig. 6C). Similarly, there was no diet-induced increase in egg production in *dilp2* mutant females (Fig. 6D). These findings suggest that the nutrient-dependent increase in IIS activity and body size are important to promote fecundity in a protein-rich context. This result aligns with findings from a previous study showing that lifetime fecundity was significantly lower in *dilp2* mutants raised in a yeast-rich diet (Grönke et al., 2010). To extend our findings beyond *dilp* genes, we next examined fecundity in females with an RNAi-mediated reduction in *sun*. We found that the nutrient-dependent increase in egg production in *r4>UAS-sun-RNAi* females was eliminated, in contrast to the robust diet-induced increase in fecundity in *r4>+* and *+>UAS-sun-RNAi* control females (Fig. 6E). Together, this data suggests that *dilp2* and fat body-derived *sun* play a role in maximizing IIS activity and body size to promote egg production in a protein-rich context.

In males, which have a reduced ability to augment body size in response to a protein-rich diet, we also investigated the relationship between nutrient content, body size, and fertility. When we compared fertility in *w^1118^* males reared on 1Y compared with males raised on 2Y, we found no significant difference in the number of offspring produced (Fig. 6F). Thus, neither male body size nor fertility were enhanced by rearing flies in a protein-rich environment. Given that previous studies suggest that a larger body size in males promotes reproductive success (Ewing, 1961; Partridge et al., 1987; Partridge & Farquhar, 1983), we next asked whether genetic manipulations that augment male body size also increased fertility. One way to augment male body size in 1Y is heterozygous loss of *phosphatase and tensin homolog* (*pten*, FBgn0026379; *pten^2L100^/+*) (Fig. S6B). Interestingly, fertility was not significantly higher in *pten^2L100^/*+ males compared with *w^1118^* controls raised in 1Y (Fig. 6F), suggesting that a larger body size does not always augment fertility in males. Similarly, when we measured fertility in *r4>UAS-sun* males, which are larger than control males (Fig. S13B), fertility was not significantly different from *r4>+* and *+>UAS-sun* control males (Fig. 6G). Thus, in males the relationship between body size and fertility is less robust than in females, as genetic manipulations that increase body size do not augment fertility. Interestingly, when we examined fertility in *pten^2L100^/*+ and *r4>UAS-sun* males in 2Y, fertility was significantly increased in *pten^2L100^/*+ males compared with genotype-matched controls cultured in 1Y (Fig. 6F), an observation we did not repeat in *r4>UAS-sun* males (Fig. 6G). Ultimately, this less robust and more complex relationship between body size and fertility in males suggests a possible explanation for their decreased nutrient-dependent body size plasticity compared with females.

## DISCUSSION

In many animals, body size plasticity in response to environmental factors such as nutrition differs between the sexes (Fairbairn, 1997). While past studies have identified mechanisms underlying nutrient-dependent growth in a mixed-sex population, and revealed factors that promote sex-specific growth in a single nutritional context, the mechanisms underlying the sex difference in nutrient-dependent body size plasticity remain unknown. In this study, we showed that females have higher phenotypic plasticity compared with males when reared on a protein-rich diet, and elucidated the molecular mechanisms underlying the sex difference in nutrient-dependent body size plasticity in this context. Our data suggests a model in which high levels of dietary protein augment female body size by stimulating an increase in IIS activity, where we identified a requirement for *dilp2* and *sun* in promoting this nutrient-dependent increase in IIS activity. Importantly, we discovered *tra* as the factor responsible for stimulating *sun* mRNA levels and IIS activity, identifying a novel role for sex determination gene *tra* in regulating nutrient-dependent body size plasticity. Together, our findings reveal one mechanism underlying the sex difference in nutrient-dependent body size plasticity.

One interesting finding from our study was the identification of a sex difference in nutrient-dependent changes to IIS activity. In females raised on a protein-rich diet, there was a nutrient-dependent upregulation of IIS activity. In males, this diet-induced increase in IIS activity was not observed. This reveals a previously unrecognized sex difference in the coupling between IIS activity and dietary protein: females tightly couple nutrient input with IIS activity across a wide protein concentration range, whereas the close coordination between dietary protein and IIS activity in males was lost in a protein-rich context. Our data shows that this sex difference in nutrient-dependent changes to IIS activity during development is physiologically significant, as it supports an increased rate of growth and consequently larger body size in females but not in males raised on a protein-rich diet. In future studies, it will be important to determine whether the sex difference in coupling between nutrients and IIS activity exist in other contexts. For example, previous studies on the extension of life span by dietary restriction have shown that male and female flies differ in the concentration of nutrients that produces the maximum life span extension, and in the magnitude of life span extension produced by dietary restriction (Magwere et al., 2004; Regan et al., 2016). Similar sex-specific effects of dietary restriction and reduced IIS on life span have also been observed in mice (Holzenberger et al., 2003; Kane et al., 2018; reviewed in Regan & Partridge, 2013; Selman et al., 2008) and humans (van Heemst et al., 2005). Future studies will be needed to determine whether a male-female difference in coupling between nutrients and IIS activity similarly explain these sex-specific life span responses to dietary restriction. Indeed, given that sex differences have been reported in the risk of developing diseases associated with overnutrition and dysregulation of IIS activity such as obesity and type 2 diabetes (Kautzky-Willer et al., 2016; Mauvais-Jarvis, 2018; Tramunt et al., 2020), more detailed knowledge of the male-female difference in coupling between nutrients and IIS activity in other models may provide insights into this sex-biased risk of disease.

In addition to revealing a sex difference in the nutrient-dependent upregulation of IIS activity, our data identified a female-specific requirement for *dilp2* and *sun* in mediating the diet-induced increase in IIS activity in a protein-rich context. While previous studies have shown that both *dilp2* and *sun* positively regulate body size (Ikeya et al., 2002; Grönke et al., 2010; Delanoue et al 2016), we describe new sex-specific roles for *dilp2* and *sun* in mediating nutrient-dependent phenotypic plasticity. Elegant studies have shown that *sun* is a secreted factor that stimulates dILP2 release from the IPCs (Delanoue et al., 2016). Together with our data, this suggests a model in which the sex difference in nutrient-dependent body size plasticity is due to the diet-induced upregulation of *sun* in females and not males. Higher *sun* mRNA levels enhance dILP2 secretion to promote IIS activity and increase female body size in a protein-rich context. This model aligns well with findings from two previous studies on dILP2 secretion in male and female larvae. The first study, which raised larvae on a protein-rich diet equivalent to the 2Y diet, found increased dILP2 secretion in females compared to males (Rideout et al., 2015). The second study, which raised larvae on a diet equivalent to the 1Y diet, found no sex difference in dILP2 secretion and no effects of *dilp2* loss on body size (Sawala & Gould, 2017). Thus, while these previous studies differed in their initial findings on a sex difference in dILP2 secretion, our data reconcile these minor differences by identifying context-dependent effects of *dilp2* on body size. Future studies will need to determine whether these sex-specific and context-dependent effects of *dilp2* are observed in other phenotypes regulated by *dilp2* and other *dilp* genes. For example, flies carrying mutations in *dilp* genes show changes to aging, metabolism, sleep, and immunity, among other phenotypes (Bai et al., 2012; Brown et al., 2020; Cong et al., 2015; Grönke et al., 2010; Liu et al., 2016; Nässel & Vanden Broeck, 2016; Okamoto et al., 2009; Okamoto & Nishimura, 2015; Post et al., 2018, 2019; Slaidina et al., 2009; Stafford et al., 2012; Zhang et al., 2009; Bai et al., 2012; Brogiolo et al., 2001; Brown et al., 2020; Cognigni et al., 2011; Cong et al., 2015; Grönke et al., 2010; Linneweber et al., 2014; Liu et al., 2016; Okamoto et al., 2009; Post et al., 2018, 2019; Semaniuk et al., 2018; Slaidina et al., 2009; Stafford et al., 2012; Suzawa et al., 2019; Ugrankar et al., 2018; Zhang et al., 2009). Further, it will be interesting to determine whether the sex-specific regulation of *sun* is observed in any other contexts, and whether it will influence sex differences in phenotypes associated with altered IIS activity, such as life span.

While our findings on *sun* and *dilp2* provide mechanistic insight into the molecular basis for the male-female difference in phenotypic plasticity in response to a protein-rich diet, a key finding from our study was the identification of sex determination gene *tra* as the factor that confers plasticity to females. Normally, nutrient-dependent body size plasticity is higher in females than in males in a protein-rich context. In females lacking a functional Tra protein, however, this increased nutrient-dependent body size plasticity was abolished. In males, which normally lack a functional Tra protein, ectopic Tra expression conferred increased nutrient-dependent body size plasticity. A previous study showed that on the 2Y diet Tra promotes dILP2 secretion (Rideout et al., 2015); however, our current study extends this finding by identifying *sun* as one link between Tra and dILP2. Further, by demonstrating that Tra’s regulation of IIS activity and body size is context-dependent, we identify a previously unrecognized role for Tra in regulating nutrient-dependent body size plasticity. This new role for *tra* also accounts for minor differences between previous studies on the effects of *tra* on growth during development (Mathews et al., 2017; Rideout et al., 2015; Sawala & Gould, 2017). While we extend these previous findings by showing that Tra confers nutrient-dependent body size plasticity via sex-specific regulation of *sun* mRNA and IIS activity, it remains unclear how Tra regulates *sun* mRNA levels in response to dietary protein. Future studies will need to examine the basis for this sex-specific regulation, as recent studies have expanded the number of Tra-regulated genes beyond its canonical targets genes *fruitless* (*fru*; FBgn0004652) and *doublesex* (*dsx*; FBgn0000504) (Clough et al., 2014; Hudry et al., 2016, 2019). In addition to these mechanistic studies, it will also be critical to explore how Tra couples *sun* mRNA regulation with dietary protein intake. Studies have shown that the *tra* locus is regulated both by alternative splicing and transcription (Belote et al., 1989; Boggs et al., 1987; Grmai et al., 2018; Inoue et al., 1990; Sosnowski et al., 1989), and that the Tra protein is regulated by phosphorylation (Du et al., 1998). Our study therefore highlights the importance of additional studies on the regulation of the *tra* genomic locus and Tra protein to gain mechanistic insight into its effects on nutrient-dependent body size plasticity.

While the main outcome of our work was to reveal the molecular mechanisms that regulate the sex difference in nutrient-dependent body size plasticity, we also provide some insight into how genes that contribute to nutrient-dependent body size plasticity affect female fecundity and male fertility. Our findings align well with previous studies demonstrating that increased nutrient availability during development and a larger female body size confers increased ovariole number and fertility (Green & Extavour, 2014; Mendes & Mirth, 2016; Robertson, 1957a, 1957b), as females lacking either *dilp2* or fat body-derived *sun* were unable to augment egg production in a protein-rich context. Given that previous studies demonstrate IIS activity influences germline stem cells in the ovary in adult flies (Hsu et al., 2008; Hsu & Drummond-Barbosa, 2009; Kao et al., 2015; LaFever & Drummond-Barbosa, 2005; Lin & Hsu, 2020; Su et al., 2018), there is a clear reproductive benefit that arises from the tight coupling between nutrient availability, IIS activity, and body size in females. In males, however, the relationship between fertility and body size remains less clear. While larger males are more reproductively successful both in the wild and in laboratory conditions (Ewing, 1961; Partridge & Farquhar, 1983), other studies revealed that medium-sized males were more fertile than both larger and smaller males (Lefranc & Bundgaard, 2000). Given that our study revealed no significant increase in the number of progeny produced by larger males, the fertility benefits that accompany a larger body size in males may be context-dependent. For example, studies have shown that a larger body size increases the ability of males to outcompete smaller males (Flatt, 2020; Partridge et al., 1987; Partridge & Farquhar, 1983). Thus, in crowded situations, a larger body size may provide significant fertility gains. On the other hand, in conditions where nutrients are limiting, an imbalance in the allocation of energy from food to growth rather than to reproduction may decrease fertility (Bass et al., 2007; Camus et al., 2017; Jensen et al., 2015; Wood et al., 2018). Future studies will therefore be needed to resolve the relationship between body size and fertility in males, as this will suggest the ultimate reason(s) for the sex difference in nutrient-dependent body size plasticity.

## MATERIALS AND METHODS

### Data Availability

Raw values for all data collected and displayed in this manuscript are available in S2 Table.

### Fly husbandry

Larvae were raised at a density of 50 animals per 10 ml food at 25°C (recipes in S3 Table), collected as indicated in figure legends, and sexed by gonad size. When gonad size could not be used to determine sex (*e.g*., *tra* mutants, *da-GAL4>UAS-tra^F^*), chromosomal females were identified by the presence of an X-linked GFP. Adult flies were maintained at a density of 20 flies per vial in single-sex groups.

### Fly strains

The following fly strains from the Bloomington *Drosophila* Stock Center were used: *Canton-S* (#64349), *w^1118^* (#3605), *tra^1^ (#675), Df(3L)st-j7 (#5416), srl^1^* (#14965), *InR^E19^* (#9646), TRiP control (#36303) *UAS-ilp2-RNAi* (#32475), *UAS-upd2-RNAi* (#33949)*, UAS-tra^F^* (#4590), *da-GAL4* (ubiquitous), *r4-GAL4* (fat body), *cg-GAL4* (fat body). The following fly strains from the Vienna *Drosophila* Resource Center were used in this study: *UAS-sun-RNAi* (GD23685)*, UAS-Gbp1-RNAi* (KK108755) *UAS-Gbp2-RNAi* (GD16696), *UAS-CCHa2-RNAi* (KK102257). Additional fly strains include: *dilp2, pten^2L100^, UAS-sun, tGPH (GFP-PH).* All genotypes used in the manuscript are listed in S4 Table.

### Body size

Pupal volume was measured in pupae sexed by gonad size as previously described (Delanoue et al., 2010; Marshall et al., 2012; Rideout et al., 2012, 2015). For adult weight, 5-day-old virgin male and female flies were weighed in groups of 10 in 1.5 ml microcentrifuge tubes on an analytical balance. Wing length was measured as previously described (Garelli et al., 2012).

### Developmental timing

Larvae were placed into the experimental diet ±2 hr post-hatching, and sexed using gonad size. Percent pupation was calculated by comparing the number of pupae at 12 hr intervals to the total larvae in the vial.

### Feeding behavior

Feeding behavior was quantified in sexed larvae by counting mouth hook contractions for 30 sec.

### RNA extraction and cDNA synthesis

One biological replicate represents ten larvae frozen on dry ice and stored at −80°C. Each experiment contained 3-4 biological replicates per sex, per genotype, and per diet, and each experiment was repeated twice. RNA was extracted using Trizol (Thermo Fisher Scientific; 15596018) according to manufacturer’s instructions, as previously described (Marshall et al., 2012; Rideout et al., 2012, 2015; Wat et al., 2020). cDNA synthesis was performed using the QuantiTect Reverse Transcription Kit according to manufacturer’s instructions (Qiagen; 205314).

### Quantitative real-time PCR (qPCR)

qPCR was performed as previously described (Rideout et al., 2012, 2015; Wat et al., 2020). A complete primer list is available in S5 Table.

### Fecundity and fertility

For female fecundity, single 6-day-old virgin female flies raised as indicated were crossed to three age-matched *CS* virgin males for a 24 hr mating period. Flies were transferred to fresh food vials with blue 2Y food to lay eggs. The number of eggs laid over 24 hr was quantified. For male fertility, single 6-day-old virgin males were paired with three 6-day-old virgin *CS* females to mate, and females were allowed to lay eggs for 24 hr. The number of progeny was quantified by counting viable pupae.

### Microscopy

GFP-PH larvae were picked into 1Y or 2Y food. Larvae were dissected 108 hr after egg laying (AEL) and inverted carcasses were fixed for 30 minutes in 4% paraformaldehyde in phosphate buffered saline (PBS) at room temperature. Carcasses were rinsed twice with PBS, once in 0.1% Triton-X in PBS (PBST) for 5 minutes, then incubated with Hoechst (5 μg/mL, Life Technologies H3570), LipidTOX Red (1:100, Thermo Fisher Scientific H34476), and phalloidin fluor 647 (1:1000, Abcam ab176759) in PBST for 40 min. The stained carcasses were washed with PBS and mounted in SlowFade Diamond (Thermo Fisher Scientific S36972). Images were acquired with a Leica SP5 (20X). Mean GFP intensity was quantified at the cell surface (marked by phalloidin) and in the cytoplasm using Fiji (Schindelin et al., 2012). Three cells per fat body were measured, and at least five fat bodies per sex and per diet were measured.

### Statistics and data presentation

Statistical analyses and data presentation were carried out using Prism GraphPad 6 (GraphPad Prism version 6.0.0 for Mac OS X,. Statistical tests are indicated in figure legends and all *p*-values are listed in S1 Table.

## ACKNOWLEDGEMENTS

We would like to thank Drs. Renald Delanoue and William Ja for the *UAS-stunted* strain, Dr. Bruce Edgar for the GFP-PH reporter (tGPH), and Dr. Linda Partridge for sharing the *dilp2* mutant strain. Stocks obtained from the Bloomington *Drosophila* Stock Center (NIH P40OD018537) were used in this study. We thank the TRiP at Harvard Medical School (NIH/NIGMS R01-GM084947) for providing transgenic RNAi fly stocks and/or plasmid vectors used in this study. Transgenic fly stocks and/or plasmids were also obtained from the Vienna *Drosophila* Resource Center (VDRC, www.vdrc.at). We acknowledge critical resources and information provided by FlyBase (Thurmond et al., 2018); FlyBase is supported by a grant from the National Human Genome Research Institute at the U.S. National Institutes of Health (<u>U41 HG000739</u>) and by the British Medical Research Council (<u>MR/N030117/1</u>). Funding for this study was provided by grants to EJR from the Canadian Institutes for Health Research (PJT-153072), Natural Sciences and Engineering Research Council of Canada (NSERC, RGPIN-2016-04249), Michael Smith Foundation for Health Research (16876), and the Canadian Foundation for Innovation (JELF-34879), and to IMA from the European Research Council (ERCAdG787470). JWM was supported by a 4-year CELL Fellowship from UBC, LWW was supported by a British Columbia Graduate Scholarship Award, ZS was supported by an NSERC Undergraduate Student Research Award, and BH was supported by an European Molecular Biology Organization Fellowship (aALTF782-2015).

## SUPPLEMENTAL FIGURES

**Figure S1.**
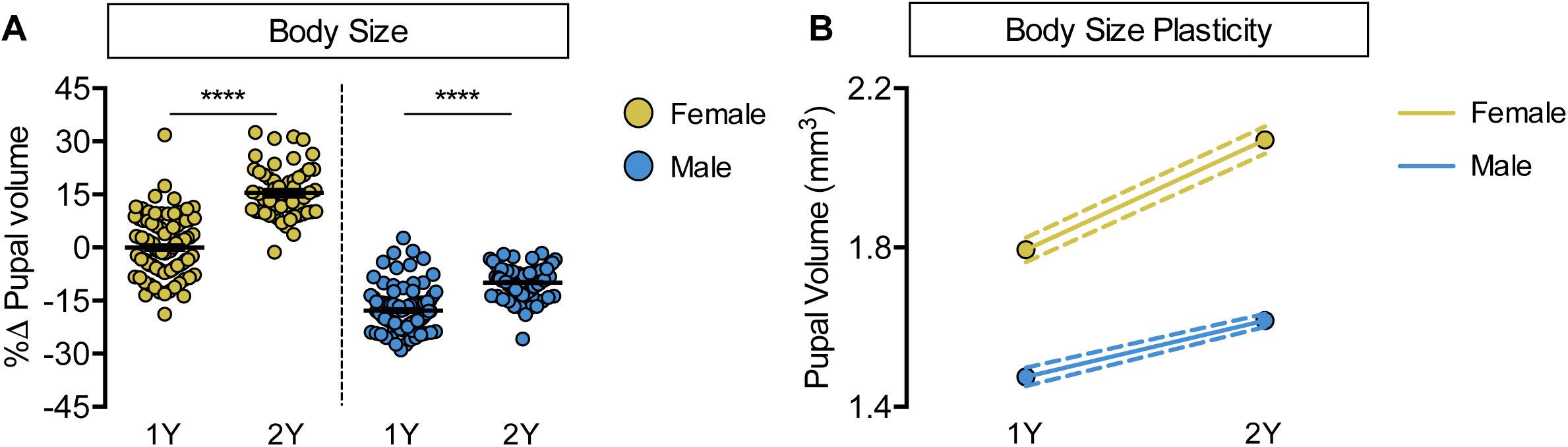
Increased nutrient-dependent body size plasticity in *Canton-S* females. (A) Pupal volume was significantly higher in both *Canton-S* (*CS*) females and males reared on a protein-rich diet (2Y) compared with genotype-matched females and males cultured on a diet containing half the protein concentration (1Y) (*p*<0.0001 for both sexes; two-way ANOVA followed by Tukey HSD test); however, the magnitude of the nutrient-dependent increase in pupal volume was higher in females (sex:diet interaction *p*<0.0001; two-way ANOVA followed by Tukey HSD test). (B) Reaction norms for pupal volume in response to changes in yeast quantity in *CS* females and males, plotted using the data in panel A. n = 57-95 pupae. For body size plasticity graphs, filled circles indicate mean pupal volume, and dashed lines indicate 95% confidence interval. **** indicates *p*<0.0001; error bars indicate SEM.

**Figure S2.**
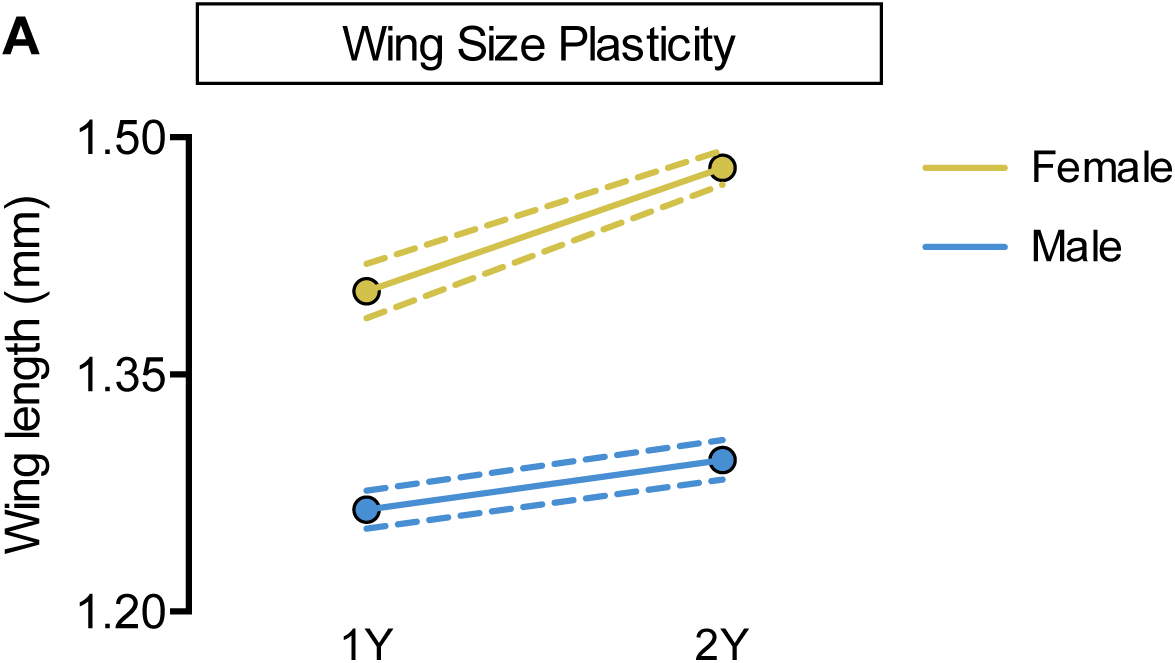
Increased nutrient-dependent plasticity in female wing size. (A) Wing length was significantly higher in both *w^1118^* females and males reared on a protein-rich diet (2Y) compared with genotype-matched females and males cultured on a diet containing half the protein content (1Y) (*p*<0.0001 and *p* = 0.0018 for females and males respectively; two-way ANOVA followed by Tukey HSD test). The magnitude of the nutrient-dependent increase in wing length was higher in females (sex:diet interaction *p* = 0.0004; two-way ANOVA followed by Tukey HSD test). n = 16-28 wings. For wing size plasticity graphs, filled circles indicate mean wing length, and dashed lines indicate 95% confidence interval.

**Figure S3.**
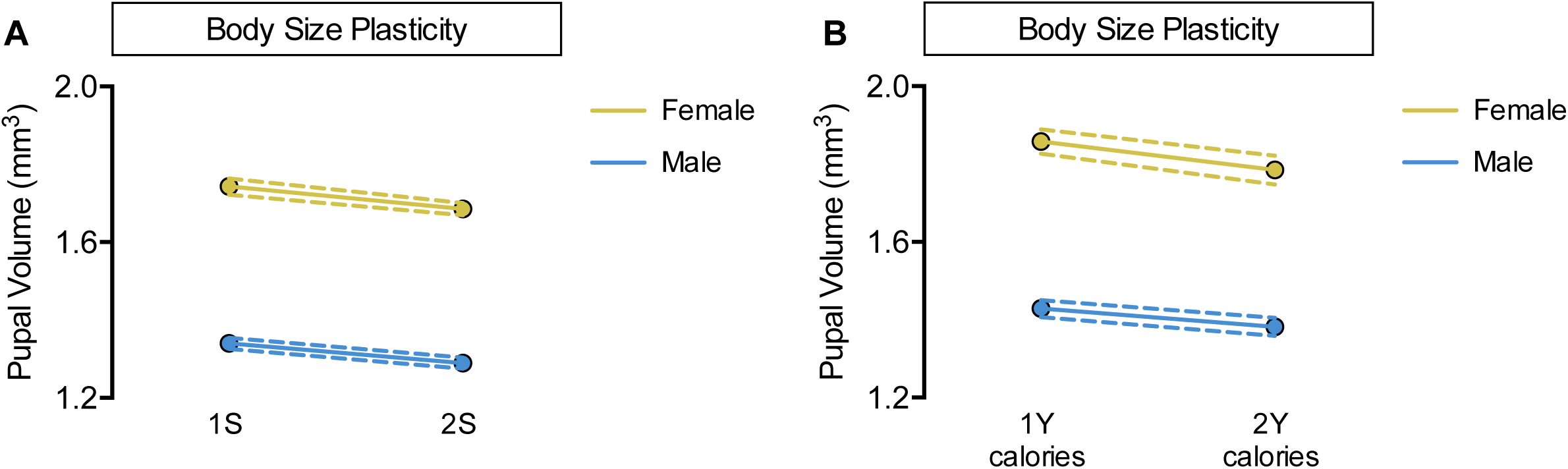
No sex-specific effect of altering dietary sugar concentration or calorie content. (A) Pupal volume was significantly decreased in both *w^1118^* females and males reared on a diet with twice the sugar (2S) compared with genotype-matched females and males cultured on a diet with the sugar content of our regular diet (1S) (*p*<0.0001 and *p* = 0.0002 for females and males respectively; two-way ANOVA followed by Tukey HSD test). The magnitude of the nutrient-dependent decrease in pupal volume was not different between females and males (sex:diet interaction *p* = 0.6536; two-way ANOVA followed by Tukey HSD test). n = 117-133 pupae. (B) While pupal volume was significantly decreased in *w^1118^* females and not males reared on a 2Y calorie-matched diet compared with genotype-matched females and males cultured on a 1Y calorie-matched diet (*p* = 0.0039 and *p* = 0.0662 respectively; two-way ANOVA followed by Tukey HSD test), there was no sex:diet interaction indicating that one sex was not more affected than the other (sex:diet interaction *p* = 0.3698; two-way ANOVA followed by Tukey HSD test). n = 44-74 pupae. For body size plasticity graphs, filled circles indicate mean pupal volume, and dashed lines indicate 95% confidence interval.

**Figure S4.**
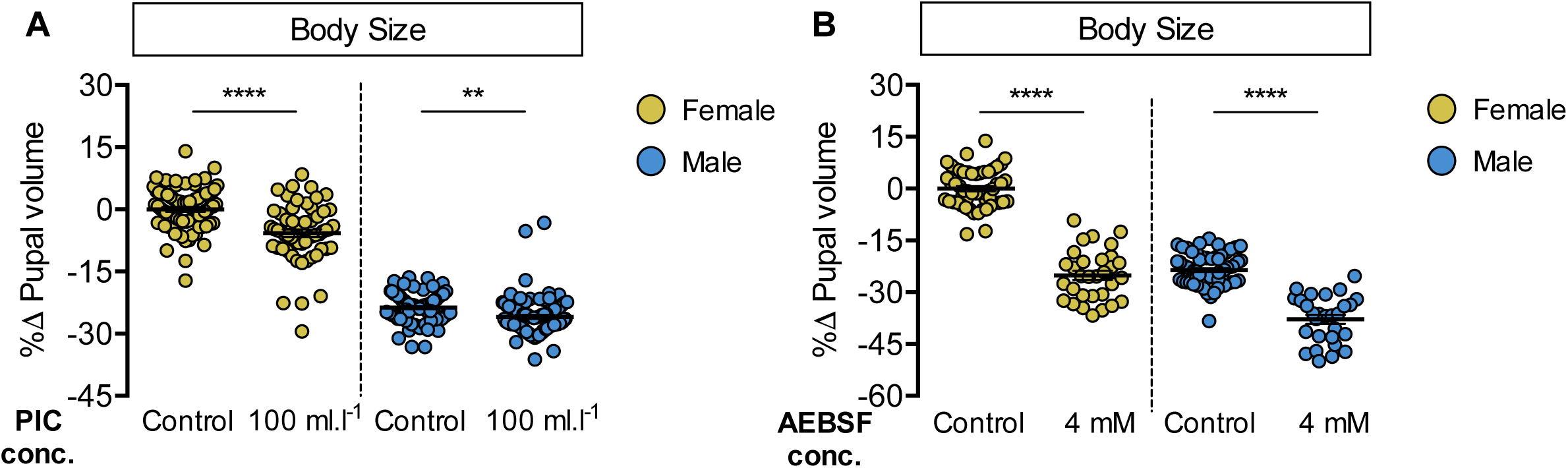
Pharmacological inhibition of protein breakdown has female-biased effects on body size. (A) Pupal volume was significantly higher in both *w^1118^* females and males reared on a protein-rich diet (2Y) compared with genotype-matched females and males cultured on 2Y containing a broad-spectrum protease inhibitor cocktail (PIC) (*p*<0.0001 and *p* = 0.0185 for females and males respectively; two-way ANOVA followed by Tukey HSD test). Importantly, the magnitude of the effect of inhibiting protein breakdown on pupal volume was higher in females (sex:treatment interaction *p* = 0.0029; two-way ANOVA followed by Tukey HSD test). n = 57-92 pupae. (B) Pupal volume was significantly higher in both *w^1118^* females and males reared on 2Y compared with genotype-matched females and males cultured on 2Y containing a serine protease-specific inhibitor 4-(2-aminoethyl)benzenesulfonyl fluoride hydrochloride (AEBSF) (*p*<0.0001 for both sexes; two-way ANOVA followed by Tukey HSD test); however, the magnitude of the effect of inhibiting protein breakdown on pupal volume was higher in females (sex:treatment interaction *p*<0.0001; two-way ANOVA followed by Tukey HSD test). n = 28-66 pupae. ** indicates *p*<0.01; **** indicates *p*<0.0001; error bars indicate SEM.

**Figure S5.**
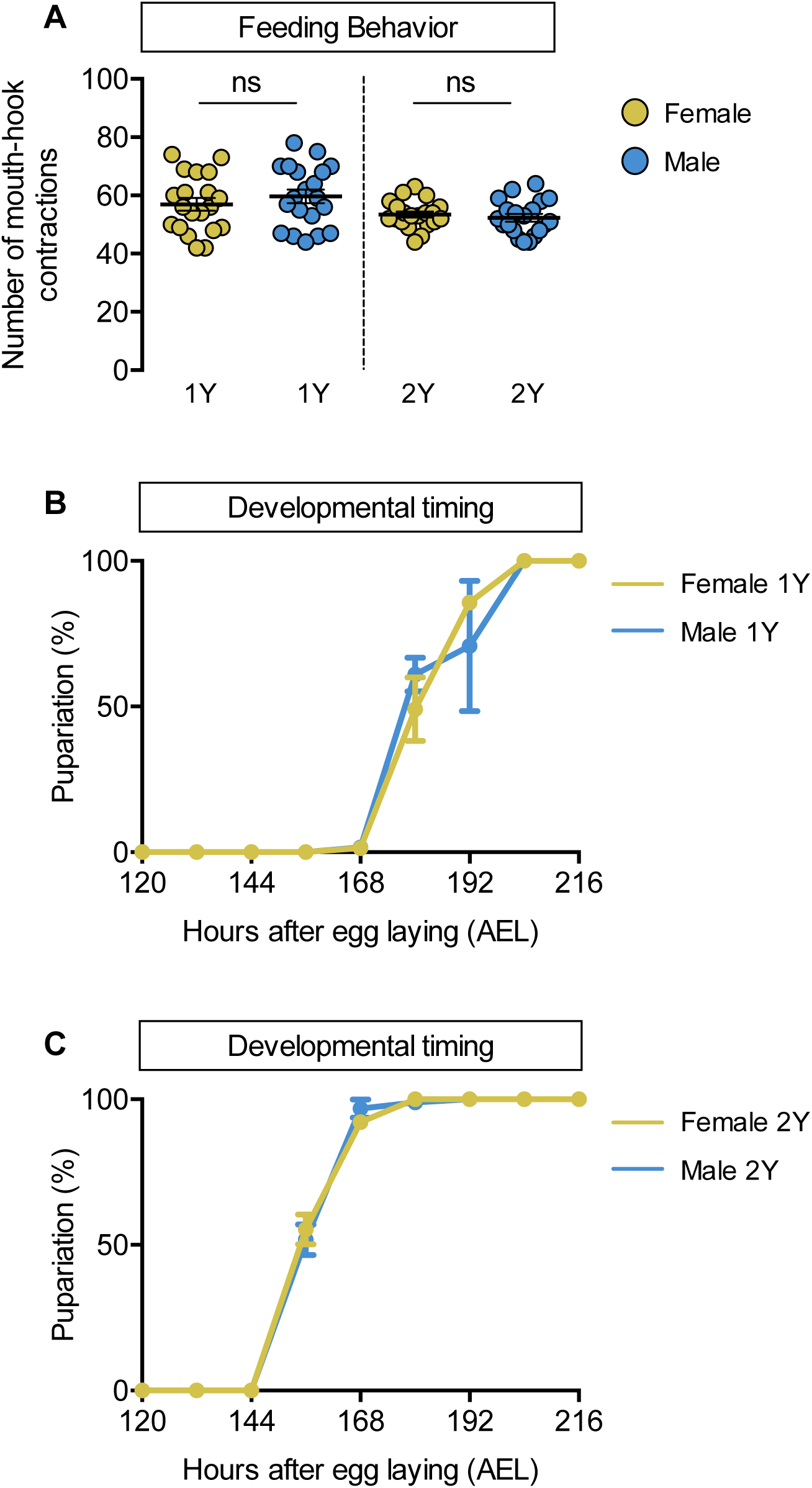
No sex difference in food intake or time to pupation. (A) There was no significant difference in mouth hook contractions between *w^1118^* control male and female larvae raised on a diet containing a widely-used protein content (1Y) (*p* = 0.3965; Student’s *t* test), or a protein-rich diet (2Y) (*p* = 0.5175; Student’s *t* test). n = 20 biological replicates. (B) There was no sex difference in the time to pupation between *w^1118^* control male and female larvae when cultured on 1Y. n = 79-93 pupae. (C) There was no sex difference in the time to pupation between *w^1118^* control male and female larvae when cultured on 2Y. n = 87-94 pupae. ns indicates not significant; error bars indicate SEM.

**Figure S6.**
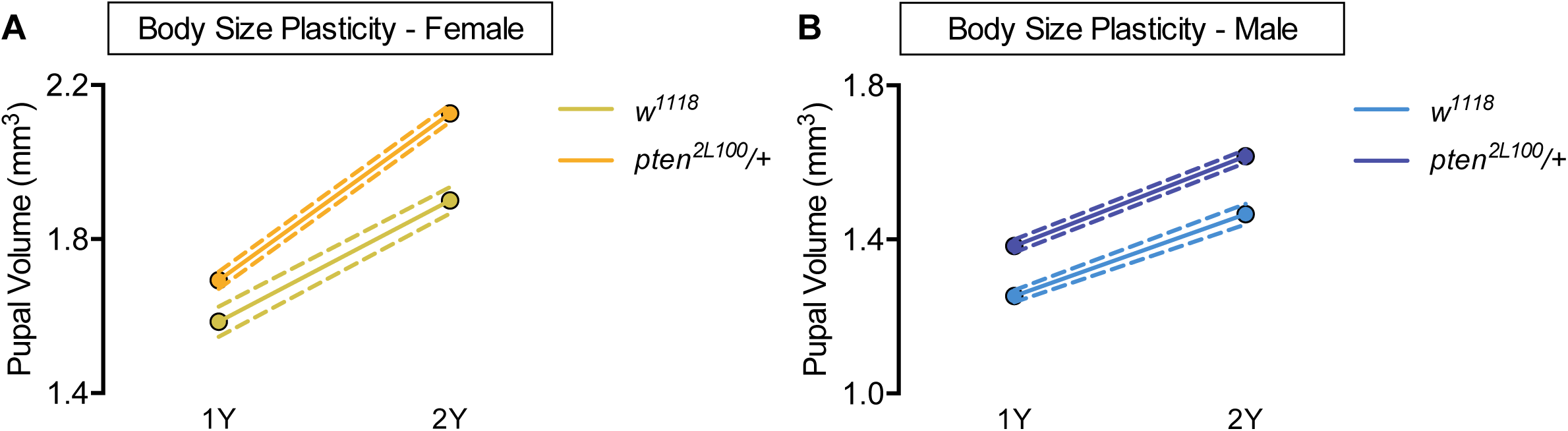
Larger body size does not confer increased body size plasticity. (A) Pupal volume was significantly higher in both *w^1118^* females and *pten^2L100^/+* females reared on a protein-rich diet (2Y) compared with genotype-matched females cultured on a diet containing half the protein content (1Y) (*p*<0.0001 for both genotypes; two-way ANOVA followed by Tukey HSD test). n = 60-89 pupae. (B) Pupal volume was significantly higher in both *w^1118^* males and *pten^2L100^/+* males reared on 2Y compared with genotype-matched males cultured on 1Y (*p*<0.0001 for both genotypes; two-way ANOVA followed by Tukey HSD test). Importantly, the magnitude of the nutrient-dependent increase in pupal volume was not different between *w^1118^* males and *pten^2L100^/+* males (genotype:diet interaction *p* = 0.3557; two-way ANOVA followed by Tukey HSD test). n = 65-88 pupae. For body size plasticity graphs, filled circles indicate mean pupal volume, and dashed lines indicate 95% confidence interval.

**Figure S7.**
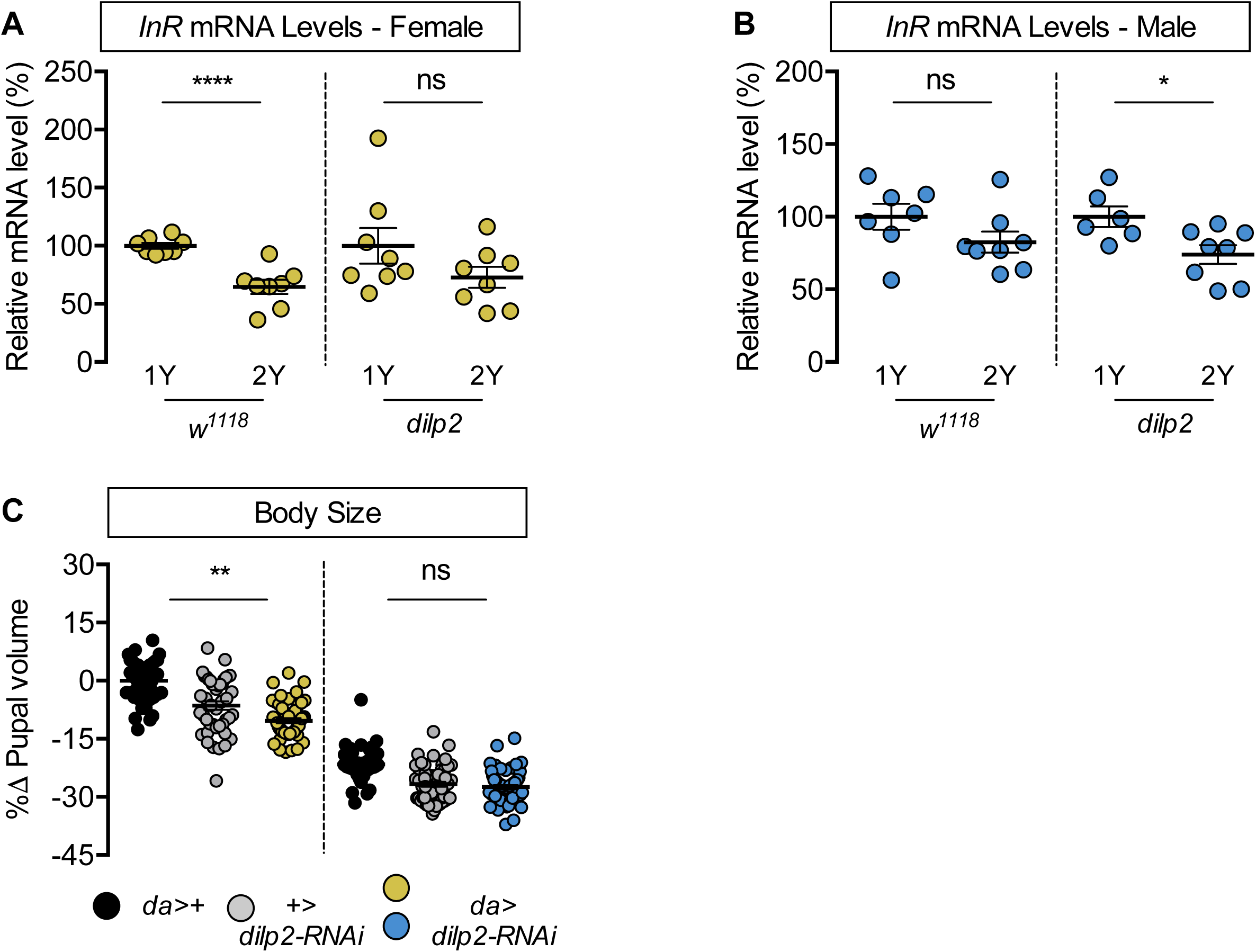
*dilp2* is required for increased nutrient-dependent body size plasticity. (A) In control *w^1118^* females, mRNA levels of *insulin receptor (InR)* were significantly lower in larvae cultured on a protein-rich diet (2Y) compared with larvae raised on a diet containing half the protein concentration (1Y) (*p*<0.0001; Student’s *t* test). In *dilp2* mutant females, there was no significant difference in *InR* mRNA levels between larvae cultured on 2Y compared with larvae raised on 1Y (*p* = 0.1472; Student’s *t* test). n = 8 biological replicates. (B) In control *w^1118^* males, mRNA levels of *InR* were not significantly lower in larvae cultured on 2Y compared with larvae raised on 1Y (*p* = 0.146; Student’s *t* test). In *dilp2* mutant males, there was a significant reduction in *InR* mRNA levels in larvae cultured on 2Y compared with larvae raised on 1Y (*p* = 0.0191; Student’s *t* test). n = 7-8 biological replicates. (C) Pupal volume was significantly reduced in females upon RNAi-mediated knockdown of *dilp2* in 2Y when compared to both control genotypes (*p*<0.0001 [*da*>+], and *p* = 0.002 [+>*UAS-dilp2-RNAi*], respectively; two-way ANOVA followed by Tukey HSD test), but not in males in 2Y (*p*<0.0001 [*da*>+], and 0.9634 [+>*UAS-dilp2-RNAi*], respectively; two-way ANOVA followed by Tukey HSD test). The magnitude of the effect of RNAi-mediated knockdown of *dilp2* on pupal volume was higher in females (sex:genotype interaction *p* = 0.003; two-way ANOVA followed by Tukey HSD test). n = 44-59 pupae. * indicates *p*<0.05, ** indicates *p*<0.01, **** indicates *p*<0.0001; ns indicates not significant; error bars indicate SEM.

**Figure S8.**
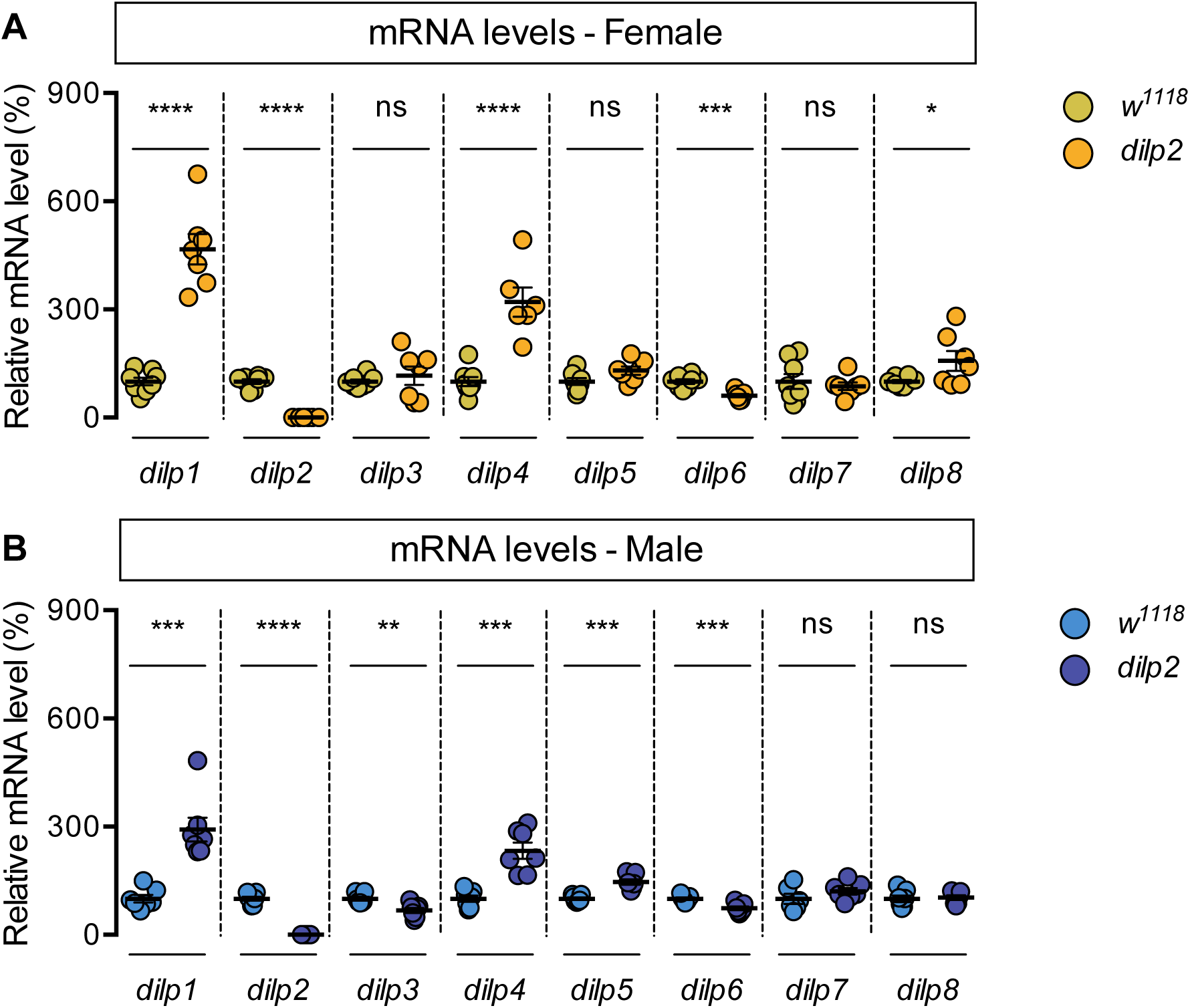
Genotype-dependent changes to *dilp* mRNA levels. (A) In *dilp2* mutant females, mRNA levels of *dilp1, dilp2, dilp4, dilp6,* and *dilp8* were significantly different from *w^1118^* control females (*p*<0.0001, <0.0001, <0.0001, 0.0003 and 0.0454, respectively; Student’s *t* test), but mRNA levels of *dilp3, dilp5,* and *dilp7* were not significantly different (*p* = 0.5142, 0.0574, and 0.605, respectively; Student’s *t* test). n = 6-8 biological replicates. (B) In *dilp2* mutant males, mRNA levels of *dilp1, dilp2, dilp3, dilp4, dilp5,* and *dilp6* were significantly different from *w^1118^* control males (*p* = 0.0001, <0.0001, 0.0034, 0.0001, 0.0001, and 0.0008, respectively; Student’s *t* test), but mRNA levels of *dilp7* and *dilp8* were not significantly different (*p* = 0.2302, and 0.7809, respectively; Student’s *t* test). n = 6-7 biological replicates. * indicates *p*<0.05, ** indicates *p*<0.01, *** indicates *p*<0.001, **** indicates *p*<0.0001; ns indicates not significant; error bars indicate SEM.

**Figure S9.**
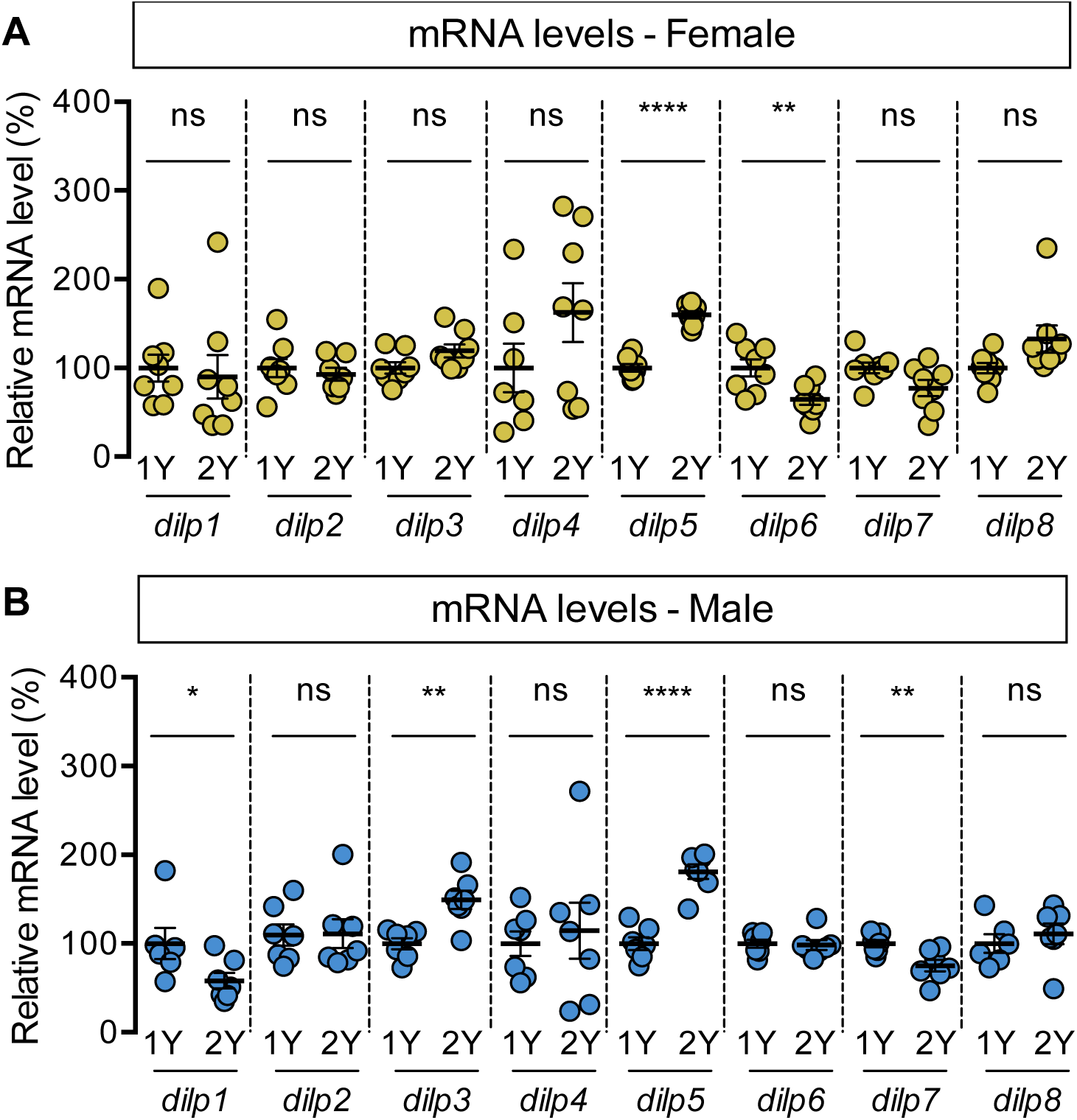
Diet-dependent changes to *dilp* mRNA levels. (A) mRNA levels of *dilp5* and *dilp6* were significantly different between females raised on a protein-rich diet (2Y) compared with female larvae cultured on a diet with half the protein concentration of 2Y (1Y) (*p*<0.0001 and 0.0079, respectively; Student’s *t* test), but mRNA levels of *dilp1, dilp2, dilp3, dilp4, dilp7, dilp8* were unchanged (*p* = 0.7337, 0.5947, 0.0672, 0.1777, 0.0562 and 0.0643, respectively; Student’s *t* test). n = 7-8 biological replicates. (B) In males cultured in 1Y, mRNA levels of *dilp1, dilp3, dilp5, dilp7* were significantly different from male larvae raised on 2Y (*p* = 0.047, 0.0014, <0.0001, and 0.0068, respectively; Student’s *t* test); mRNA levels of *dilp2, dilp4, dilp6,* and *dilp8* were unchanged (*p* = 0.9388, 0.6812, 0.8157 and 0.5054, respectively; Student’s *t* test). n = 6-7 biological replicates. * indicates *p*<0.05, ** indicates *p*<0.01, **** indicates *p*<0.0001; ns indicates not significant; error bars indicate SEM.

**Figure S10.**
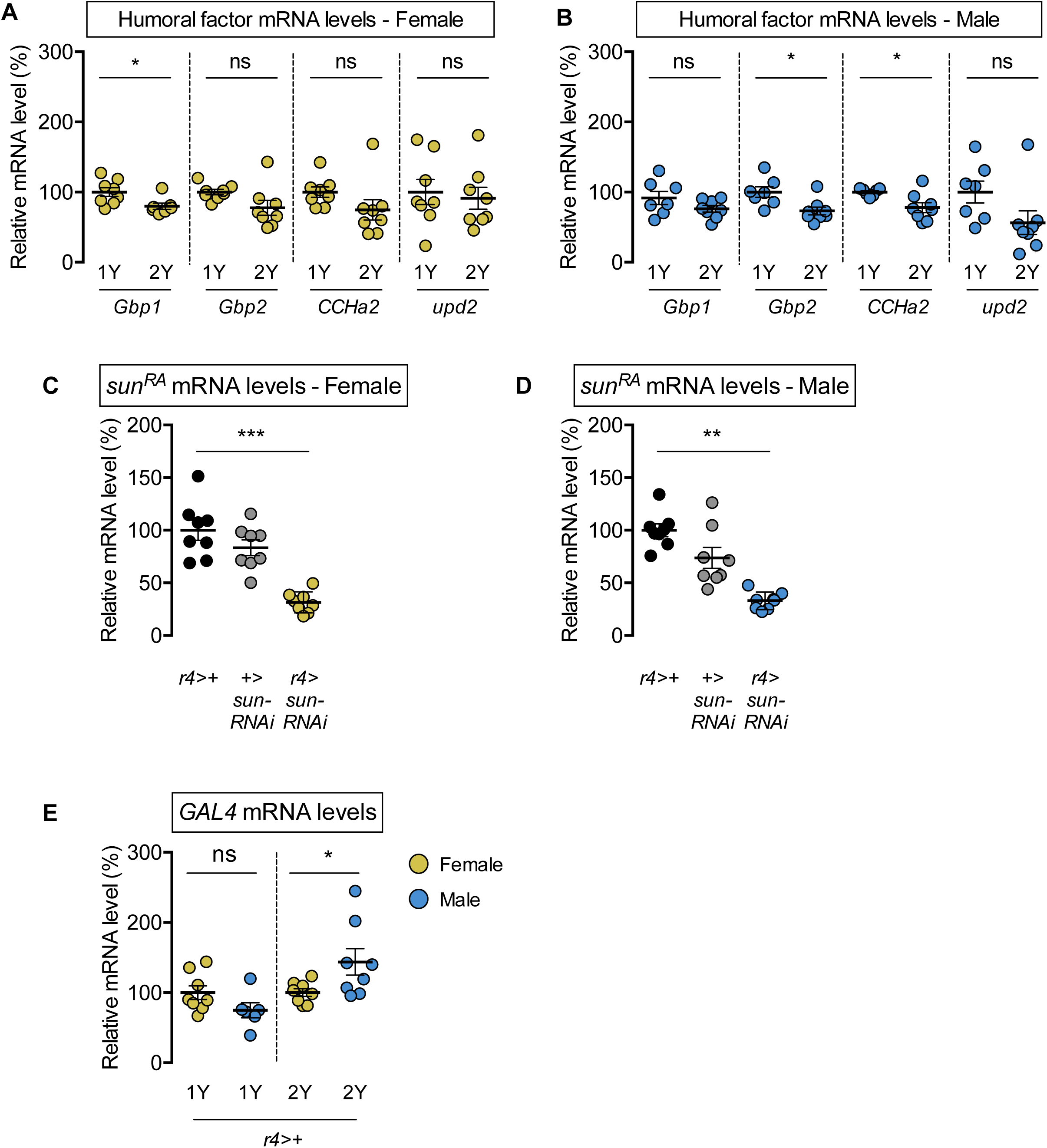
Diet-induced changes to mRNA levels of humoral factors. (A) mRNA levels of *Growth-blocking peptide 1* (*Gbp1*) were significantly different in females cultured on a protein-rich diet (2Y) compared with females raised in a diet containing half the protein concentration (1Y) (*p* = 0.0245; Student’s *t* test); however, mRNA levels of *Growth-blocking peptide 2* (*Gbp2*)*, CCHamide-2* (*CCHa2*), and *unpaired 2* (*upd2*) were not significantly different between female larvae raised on 1Y and 2Y (*p* = 0.0662, 0.1416, and 0.7171, respectively; Student’s *t* test). n = 7-8 biological replicates. (B) Levels of *Gbp1* and *upd2* were not significantly different between male larvae raised on 2Y compared with larvae reared on 1Y (*p* = 0.1487, and *p* = 0.1686, respectively; Student’s *t* test); whereas levels of *Gbp2* and *CCHa2* were significantly different between males raised in 2Y and 1Y (*p* = 0.0214, and *p* = 0.0272, respectively; Student’s *t* test). n = 7-8 biological replicates. (C) mRNA levels of *stunted* (*sun^RA^*) were significantly lower in *r4-GAL4>UAS-sun-RNAi* females compared with *r4-GAL4>+* and *+>UAS-sun-RNAi* control females (*p*<0.0001 and *p* = 0.0001, respectively; one-way ANOVA followed by Tukey HSD test). n = 8 biological replicates. (D) mRNA levels of *stunted* (*sun^RA^*) were significantly lower in *r4-GAL4>UAS-sun-RNAi* males compared with *r4-GAL4>+* and *+>UAS-sun-RNAi* control males (*p*<0.0001 and *p* = 0.0012, respectively; one-way ANOVA followed by Tukey HSD test). n = 8 biological replicates. (E) Levels of *GAL4* mRNA were not significantly different between the sexes in larvae raised in 1Y (*p* = 0.1105; Student’s *t* test), whereas *GAL4* mRNA levels were significantly higher in males in 2Y (*p* = 0.0428; Student’s *t* test). n = 6-8 biological replicates. * indicates *p*<0.05, ** indicates *p*<0.01, *** indicates *p*<0.001; ns indicates not significant; error bars indicate SEM.

**Figure S11.**
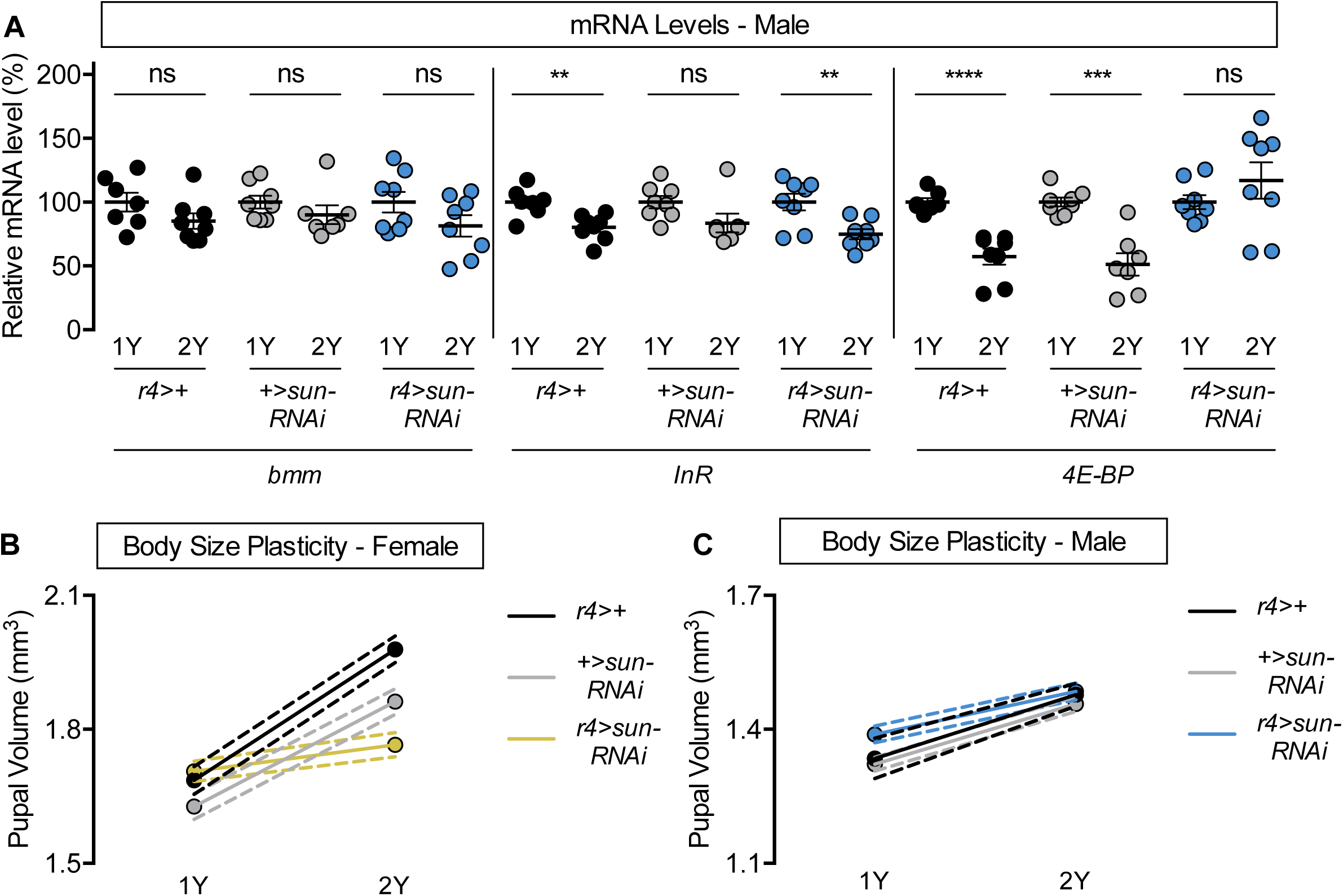
Nutrient-dependent upregulation of IIS activity and increased female body size plasticity requires *stunted* (*sun*). (A) In *r4>+*, *+>UAS-sun-RNAi* males, and *r4>UAS-sun-RNAi* males, mRNA levels of *brummer (bmm*) were not significantly different between larvae raised on a protein-rich diet (2Y) compared with larvae reared on a diet containing half the protein concentration (1Y) (*p* = 0.1445, 0.2766, and 0.1308, respectively; Student’s *t* test). In *r4>+* and *r4>UAS-sun-RNAi* males, mRNA levels of *insulin receptor* (*InR*) were significantly different in larvae between 1Y and 2Y (*p* = 0.003 and *p* = 0.0054, respectively; Student’s *t* test), but not in *+>UAS-sun-RNAi* males (*p* = 0.0745; Student’s *t* test). In *r4>+* and *+>UAS-sun-RNAi* control males, mRNA levels of *eukaryotic initiation factor 4E-binding protein* (*4E-BP*) were significantly different between larvae raised in 1Y or 2Y (*p*< 0.0001 and *p* = 0.0001, respectively; Student’s *t* test), but not in *r4>UAS-sun-RNAi* males (*p* = 0.2899; Student’s *t* test). n = 7-8 biological replicates. (B) Pupal volume was significantly higher in *r4>+*, *+>UAS-sun-RNAi*, and *r4>UAS-sun-RNAi* females reared on 2Y compared with genotype-matched females cultured on 1Y (*p*<0.0001 [*r4>+* and *+>UAS-sun-RNAi*] and *p* = 0.0367 [*r4>UAS-sun-RNAi*]; two-way ANOVA followed by Tukey HSD test). The magnitude of the nutrient-dependent increase in pupal volume was significantly lower in *r4>UAS-sun-RNAi* females (genotype:diet interaction *p*<0.0001; two-way ANOVA followed by Tukey HSD test). n = 69-80 pupae. (C) Pupal volume was significantly higher in *r4>+*, *+>UAS-sun-RNAi*, and *r4>UAS-sun-RNAi* males reared on 2Y compared with genotype-matched males cultured on 1Y (*p*<0.0001 for all genotypes; two-way ANOVA followed by Tukey HSD test). The magnitude of the nutrient-dependent increase in pupal volume was not significantly different between *r4>UAS-sun-RNAi* males and control males (genotype:diet interaction *p* = 0.0784; two-way ANOVA followed by Tukey HSD test). n = 44-80 pupae. For body size plasticity graphs, filled circles indicate mean pupal volume, and dashed lines indicate 95% confidence interval. ** indicates *p*<0.01, *** indicates *p*<0.001; **** indicates *p*<0.0001; ns indicates not significant; error bars indicate SEM.

**Figure S12.**
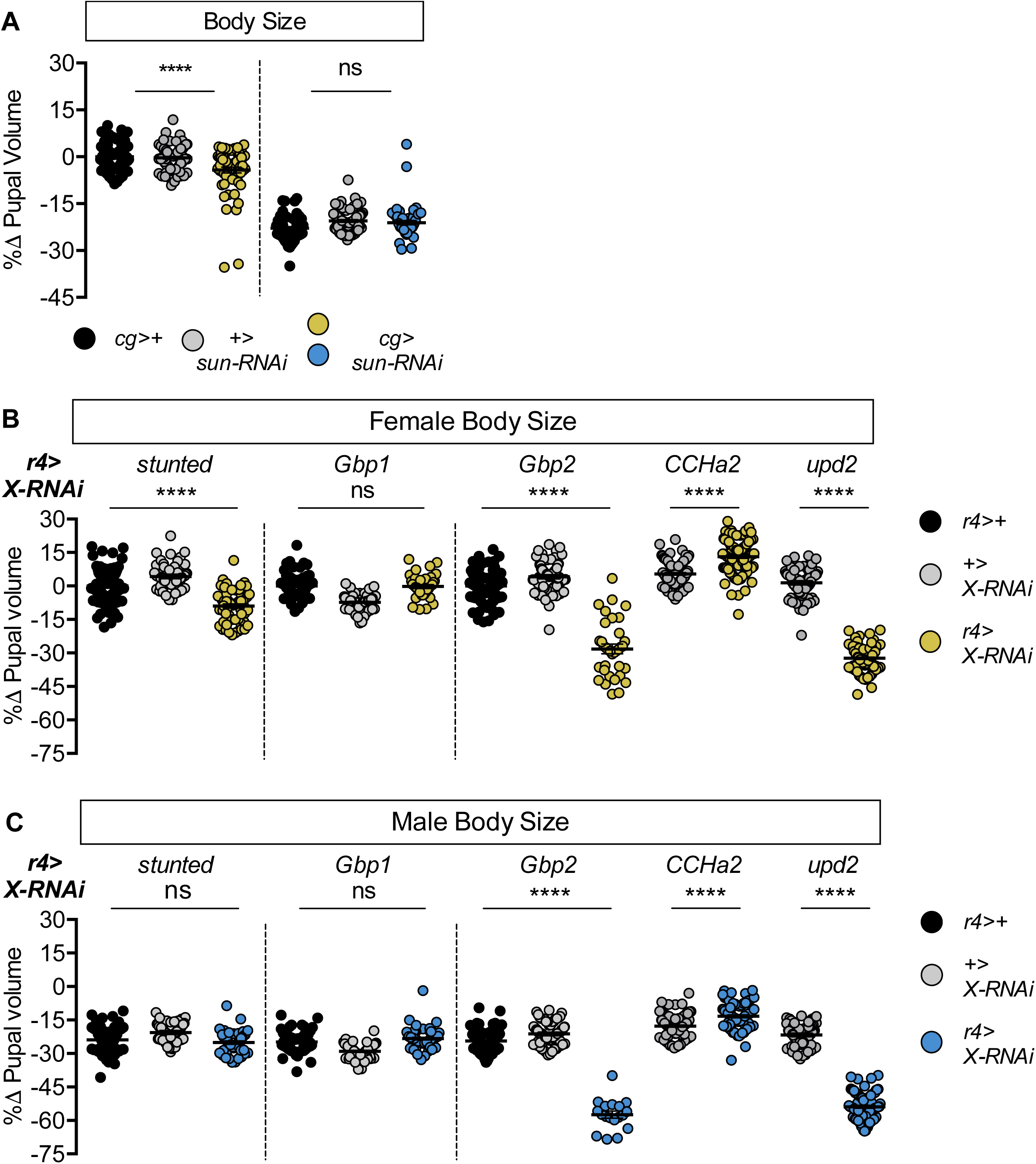
Most humoral factors have non-sex-specific effects on body size. (A) Pupal volume was significantly smaller in females with fat body-specific expression of an RNAi transgene directed against *stunted* (*sun*). Pupal volume was significantly reduced in *cg>UAS-sun-RNAi* females compared with *cg>+* and *+>UAS-sun-RNAi* control females (*p*<0.0001 for both comparisons; two-way ANOVA followed by Tukey HSD test). This decreased pupal volume was not reproduced in *cg>UAS-sun-RNAi* males compared with *cg>+* and *+>UAS-sun-RNAi* control males (*p* = 0.3657 and *p* = 0.9852, respectively; two-way ANOVA followed by Tukey HSD test). RNAi-mediated knockdown of *sun* had larger effects on pupal volume in females than in males (sex:genotype interaction *p*<0.0001; two-way ANOVA followed by Tukey HSD test). n = 54-85 pupae. (B) Pupal volume was significantly different in females with fat body-specific expression of RNAi transgenes directed against *sun*, *Growth-blocking peptide 2* (*Gbp2*), *CCHamide-2* (*CCHa2*), *unpaired 2* (*upd2*) compared with *r4>+* and *+>UAS-X-RNAi* control females (*p*<0.0001 for both comparisons [*sun*], *p*<0.0001 for both comparisons [*Gbp2*], *p*<0.0001 for both comparisons [*CCHa2*], *p*<0.0001 for both comparisons [*upd2*]; one-way ANOVA followed by Tukey HSD test); but not upon RNAi-mediated knockdown of *Growth-blocking peptide 1* (*Gbp1*) (*p* = 0.9665 and *p*<0.0001 respectively; one-way ANOVA followed by Tukey HSD test). n = 35-114 pupae. (C) Pupal volume was significantly different in males with fat body-specific expression of RNAi transgenes directed against *Gbp2*, *CCHa2*, and *upd2* compared with *r4>+* and *+>UAS-X-RNAi* control males (*p*<0.0001 for both comparisons [*Gbp2*], *p*<0.0001 for both comparisons [*CCHa2*], *p*<0.0001 for both comparisons [*upd2*]; one-way ANOVA followed by Tukey HSD test); but not reduced in males carrying RNAi transgenes directed against *sun* and *Gbp1* (*p =* 0.3513 and *p*<0.0001, respectively [*sun*]; *p =* 0.1274 and *p*<0.0001, respectively [*Gbp1*]; one-way ANOVA followed by Tukey HSD test). n = 18-100 pupae. For body size graphs, filled circles indicate pupal volume and error bars indicate SEM. **** indicates *p*<0.0001; ns indicates not significant.

**Figure S13.**
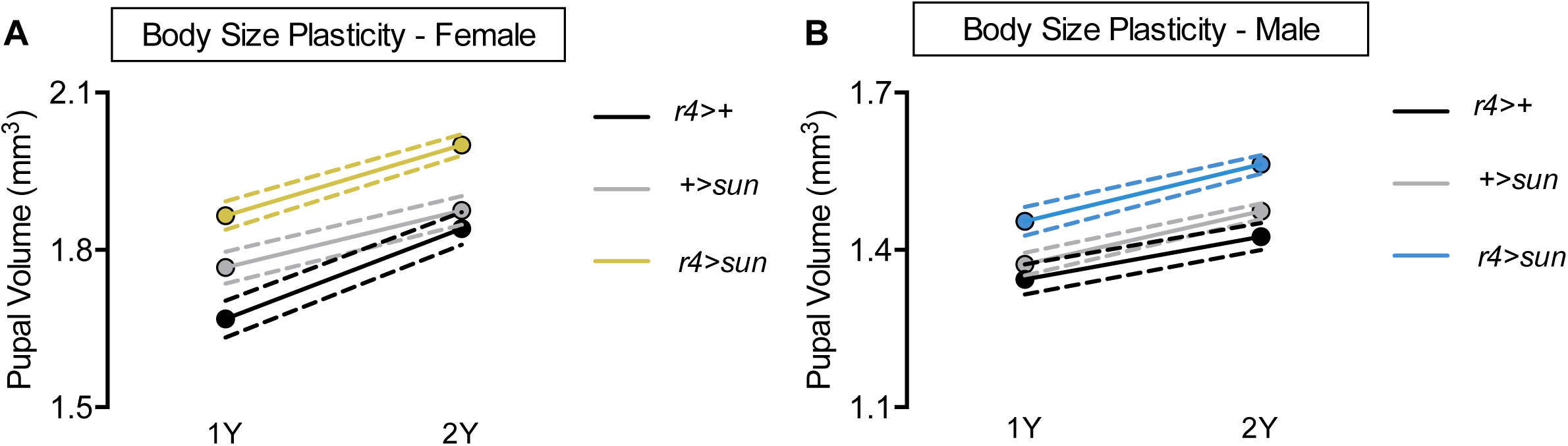
*stunted* (*sun*) overexpression augments body size but does not confer increased body size plasticity in males. (A) Pupal volume was significantly higher in *r4>+*, *+>UAS-sun*, and *r4>UAS-sun* females reared on a protein-rich diet (2Y) compared with genotype-matched females cultured on a diet containing half the protein concentration (1Y) (*p*<0.0001 for all genotypes; two-way ANOVA followed by Tukey HSD test). The magnitude of the nutrient-dependent increase in pupal volume was not significantly different between female genotypes (genotype:diet interaction *p* = 0.0895; two-way ANOVA followed by Tukey HSD test). n = 43-65 pupae. (B) Pupal volume was significantly higher in *r4>+*, *+>UAS-sun*, and *r4>UAS-sun* males reared on 2Y compared with genotype-matched males cultured on 1Y (*p*<0.0001 for all genotypes; two-way ANOVA followed by Tukey HSD test), but the magnitude of the nutrient-dependent increase in pupal volume was not different between male genotypes (genotype:diet interaction *p* = 0.4959; two-way ANOVA followed by Tukey HSD test). n = 44-67 pupae. For body size plasticity graphs, filled circles indicate mean pupal volume, and dashed lines indicate 95% confidence interval.

**Figure S14.**
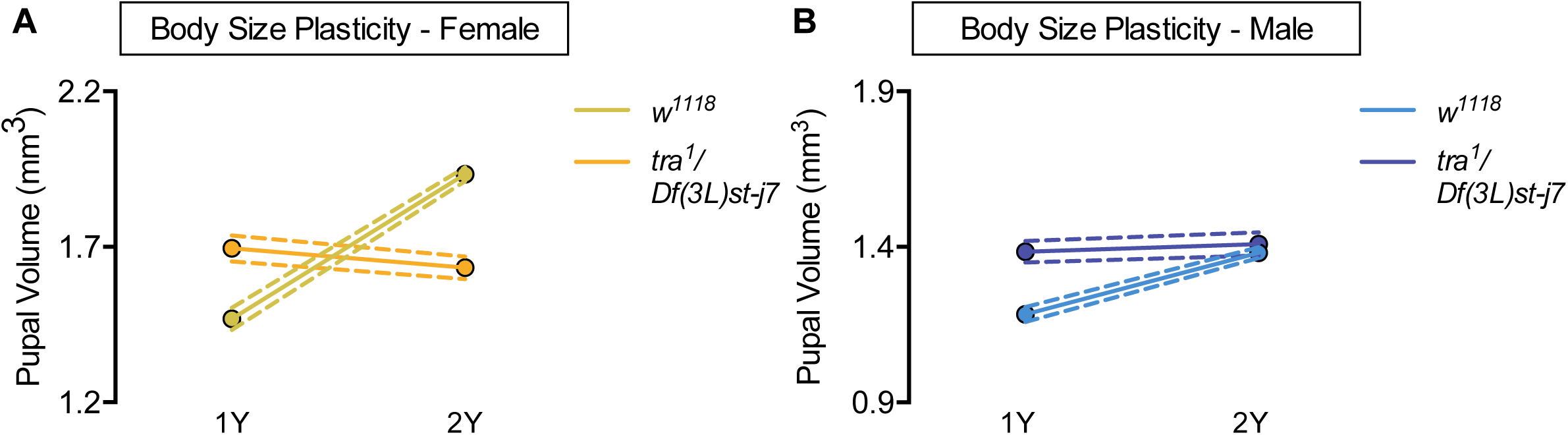
Increased nutrient-dependent body size plasticity in females requires *transformer*. (A) Pupal volume was significantly higher in *w^1118^* females reared on a protein-rich diet (2Y) compared with *w^1118^* females cultured on a diet containing half the protein concentration (1Y) (*p*<0.0001; two-way ANOVA followed by Tukey HSD test); however, this nutrient-dependent increase in pupal volume was not observed in *transformer* (*tra*) mutant females (*tra^1^/Df(3L)st-j7*) (*p* = 0.1036; two-way ANOVA followed by Tukey HSD test). The magnitude of the nutrient-dependent increase in pupal volume was lower in *tra^1^/Df(3L)st-j7* females (genotype:diet interaction *p*<0.0001). n = 39-69 pupae. (B) Pupal volume was significantly higher in *w^1118^* males (*p*<0.0001; two-way ANOVA followed by Tukey HSD test), but not in *tra^1^/Df(3L)st-j7* mutant males reared on 2Y compared with genotype-matched females cultured on 1Y (*p* = 0.6643; two-way ANOVA followed by Tukey HSD test). n = 37-65 pupae. For body size plasticity graphs, filled circles indicate mean pupal volume, and dashed lines indicate 95% confidence interval.

**Figure S15.**
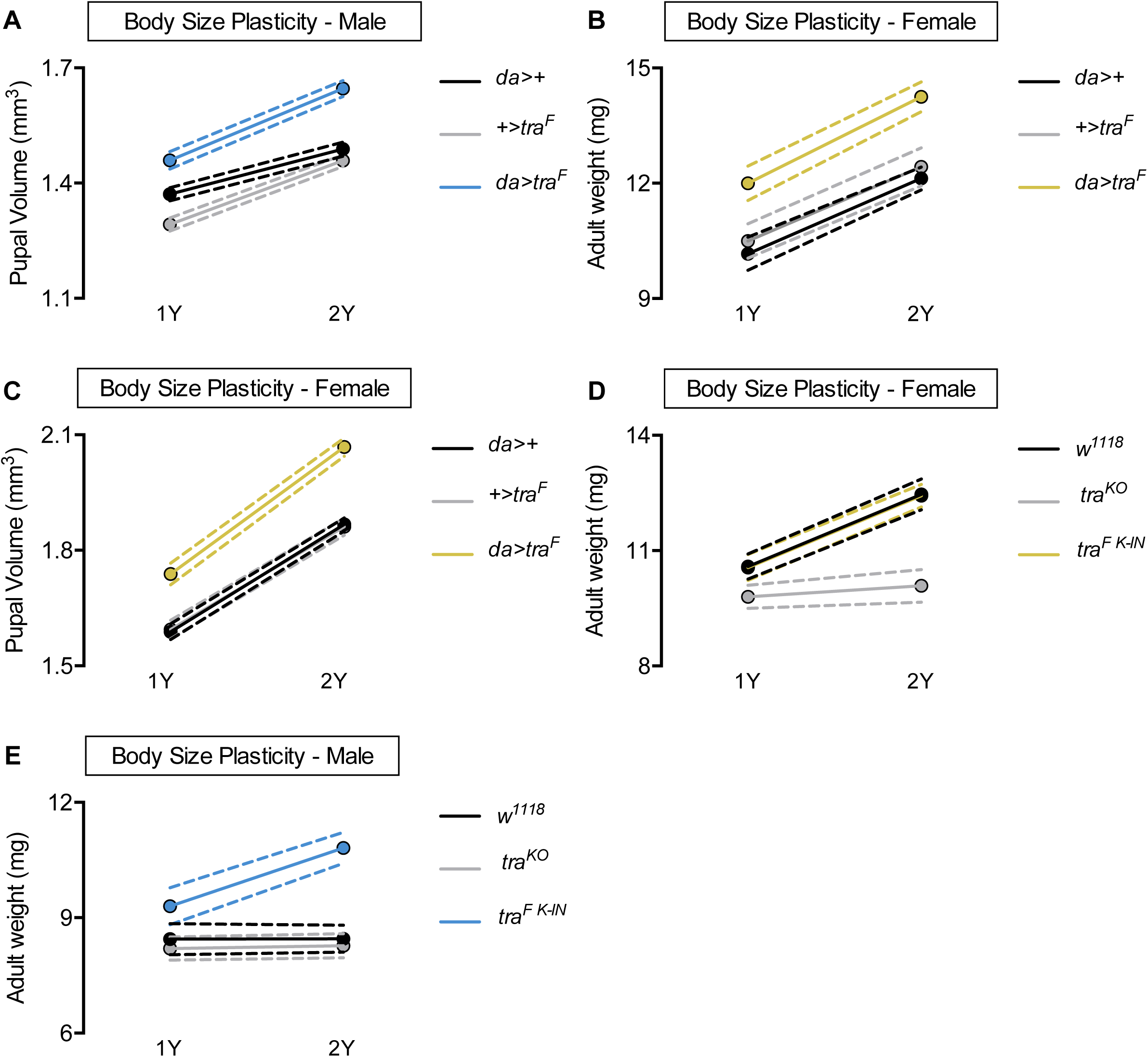
Sex determination gene *transformer* (*tra*) regulates increased nutrient-dependent body size plasticity. (A) Pupal volume was significantly higher in *da>+*, *+>UAS-tra^F^*, and *da>UAS-tra^F^* males reared on a protein-rich diet (2Y) compared with genotype-matched males cultured on a diet containing half the protein concentration (1Y) (*p*<0.0001 for all genotypes; two-way ANOVA followed by Tukey HSD test). Importantly, the magnitude of the nutrient-dependent increase in pupal volume was higher in *da>UAS-tra^F^* males (genotype:diet interaction *p* = 0.0012; two-way ANOVA followed by Tukey HSD test). n = 70-91 pupae. (B) Adult weight was significantly higher in *da>+*, *+>UAS-tra^F^*, and *da>UAS-tra^F^* females reared on 2Y compared with genotype-matched females cultured on 1Y (*p*<0.0001 for all genotypes; two-way ANOVA followed by Tukey HSD test). The magnitude of the nutrient-dependent increase in adult weight was not significantly different between *da>UAS-tra^F^* females and *da>+* and *+>UAS-tra^F^* controls (genotype:diet interaction *p* = 0.5912; two-way ANOVA followed by Tukey HSD test). n = 6-8 groups of 10 flies. (C) Pupal volume was significantly higher in *da>+*, *+>UAS-tra^F^*, and *da>UAS-tra^F^* females reared on 2Y compared with genotype-matched females cultured on 1Y (*p*<0.0001 for all genotypes; two-way ANOVA followed by Tukey HSD test). n = 68-94 pupae. (D) Adult weight was significantly higher in both *w^1118^* females, and in females with a knock-in transgene of the female isoform of *tra* (*tra^F K-IN^*), when reared on 2Y compared with 1Y (*p*<0.0001 for both genotypes; two-way ANOVA followed by Tukey HSD test). In contrast, the nutrient-dependent increase in adult weight was abolished in *tra* mutant females (*tra^KO^*) reared on 2Y compared with genotype-matched females cultured on 1Y (*p* = 0.864; two-way ANOVA followed by Tukey HSD test). Importantly, the magnitude of the nutrient-dependent increase in adult weight was significantly lower in *tra^KO^* females, which lack a functional Tra protein, than in *w^1118^* and *tra^F K-IN^* females (genotype:diet interaction *p*<0.0001; two-way ANOVA followed by Tukey HSD test). n = 10-16 groups of 10 flies. (E) Adult weight was significantly higher in *tra^F K-IN^* males, which express physiological levels of a functional Tra protein, when the males were reared on 2Y compared with genotype-matched males raised on 1Y (*p*<0.0001; two-way ANOVA followed by Tukey HSD test). In contrast, there was no significant increase in adult weight in *w^1118^* and *tra^KO^* male flies reared on 2Y compared with genotype-matched males raised on 1Y (*p*>0.9999 and *p* = 0.9996, respectively; two-way ANOVA followed by Tukey HSD test). The magnitude of the nutrient-dependent increase in adult weight was significantly higher in *tra^F K-IN^* males compared with *w^1118^* and *tra^KO^* male flies (genotype:diet interaction *p*<0.0001; two-way ANOVA followed by Tukey HSD test). n = 9-11 groups of 10 flies. For body size plasticity graphs, filled circles indicate mean pupal volume, and dashed lines indicate 95% confidence interval.

